# Low prevalence of natural *Wolbachia* in major malarial vectors *Anopheles culicifacies* s.l., and *Anopheles stephensi:* a first report

**DOI:** 10.1101/2020.11.22.393652

**Authors:** S. Gowri Sankar, T. Mowna Sundari, A. Alwin Prem Anand

**Affiliations:** ICMR-Vector Control Research Centre - Field Station, Madurai - 625002, Tamil Nadu, India; DBT - BIF Centre (Under DBT BTISNet Scheme), Lady Doak College, Madurai - 625002, Tamil Nadu, India; Department of Biotechnology, Lady Doak College, Madurai - 625002, Tamil Nadu, India

**Keywords:** *Anopheles culicifacies* s.l., *Anopheles stephensi*, *Wolbachia*, malarial vector, *16S* rRNA, endosymbiont

## Abstract

*Wolbachia* is an alpha-proteobacteria present in several arthropods. The present study focussed on the identification of *Wolbachia* in *Anopheles culicifacies* s.l., and *Anopheles stephensi*, which are wild malarial vector mosquitoes in India. A total of four hundred and eighty seven adult female mosquito samples were collected using taxonomic keys and confirmed by molecular analysis where 48.25% was *Anopheles culicifacies* s.l., and 51.75% was *Anopheles stephensi*. The presence of *Wolbachia* was identified using *16S* rRNA, *wsp* and *FtsZ* genes, where nested PCR of *16S* rRNA alone was successful and sequenced. Only seven mosquitoes (1.4%) were positive for *Wolbachia. In silico* and restriction digestion of *16S* rRNA gene product using RsaI enzyme showed that the identified *Wolbachia* belongs to supergroup B, which is further supported by phylogenetic analysis. Low prevalence rate of natural *Wolbachia* was observed in *An. culicifacies* s.l. (1.7%) and *An. stephensi* (1.2%). This is the first report on the presence of *Wolbachia* in *Anopheles culicifacies* s.l. and *Anopheles stephensi*.

## Introduction

*Wolbachia* is an intracellular alpha-proteobacteria found in a wide range of arthropods. It was first discovered by Hertig and Wolbach in 1924 considered to be the abundant endosymbiont found in invertebrates [1] and cause reproductive abnormalities (cytoplasmic incompatibility, male killing, parthenogenesis and feminization) in the host [2,3]. This endosymbiotic proteobacteria naturally infect 65% of insect species [2,4,5] including the family *Culicidae* [6,7]. So far *Wolbachia* wildtype is reported in the following mosquito species: *Culex* (*Cx*.) *pipiens* [1], *Cx. quinquefasciatus* [8], *Aedes* (*Ae*.) *albopictus* [9,10], *Ae. aegypti* [11] and few *Anopheles* species [12,13].

The natural occurrence of *Wolbachia* in *Anopheles* species has not been extensively studied. The *Anopheles* genera of *Culicidae* consists of 537 species [14], where 41 species are dominant vector species (DVS) responsible for the transmission of malaria [15]. Among the 41 DVS, 19 species/species complex were found within Asian-Pacific region [16]. As per WHO, 228 million malaria cases occurred worldwide in 2018; where India is one among the twenty countries that carries 85% of the global malarial burden [17]. In India, *An. baimaii, An. fluviatilis, An. minimus, An. sundaicus, An. culicifacies* complex species (A, B, C, D & E) and *An. stephensi* are primary vectors in transmitting malaria, with *An. culicifacies* complex and *An. stephensi* as major contributors [18]. *An. culicifaciess. s*.*l*. is widely distributed in rural, semiurban and forest areas [18-20] and, *An. stephensi* in peri-, semi- and urban areas [18,19].

In Tamil Nadu, *An. stephensi* [21-23] and *An. culicifacies* [22,24,25] are the main malaria vectors. Interestingly, few reports on natural *Wolbachia* endosymbionts in major malarial vector *Anopheles* species are reported including *An. gambiae, An. coluzzii* [12,13], *An. arabiensis* [26] and *An. moucheti* [27,28]. The present study aimed at the investigation of *Wolbachia* infection in wild mosquito population collected from different geographical locations in Tamilnadu.

## Materials and Methods

### Mosquito collection and taxonomy

Adult female mosquito samples were collected using nets and aspirators, from the 5 different locations along the foothills of the Western Ghats, Southern India (Fig.1) and identified initially by taxonomic keys [29,30] and later verified by DNA barcoding. During collection, the mosquitoes bearing ectoparasites mites has been discarded and not been used in any experiments. In the field, each samples was washed twice in 95% ethanol to avoid external contamination [31], and transferred in vial containing 95% ethanol for preservation [32] and brought to the lab for further experiments.

**Fig.1:**
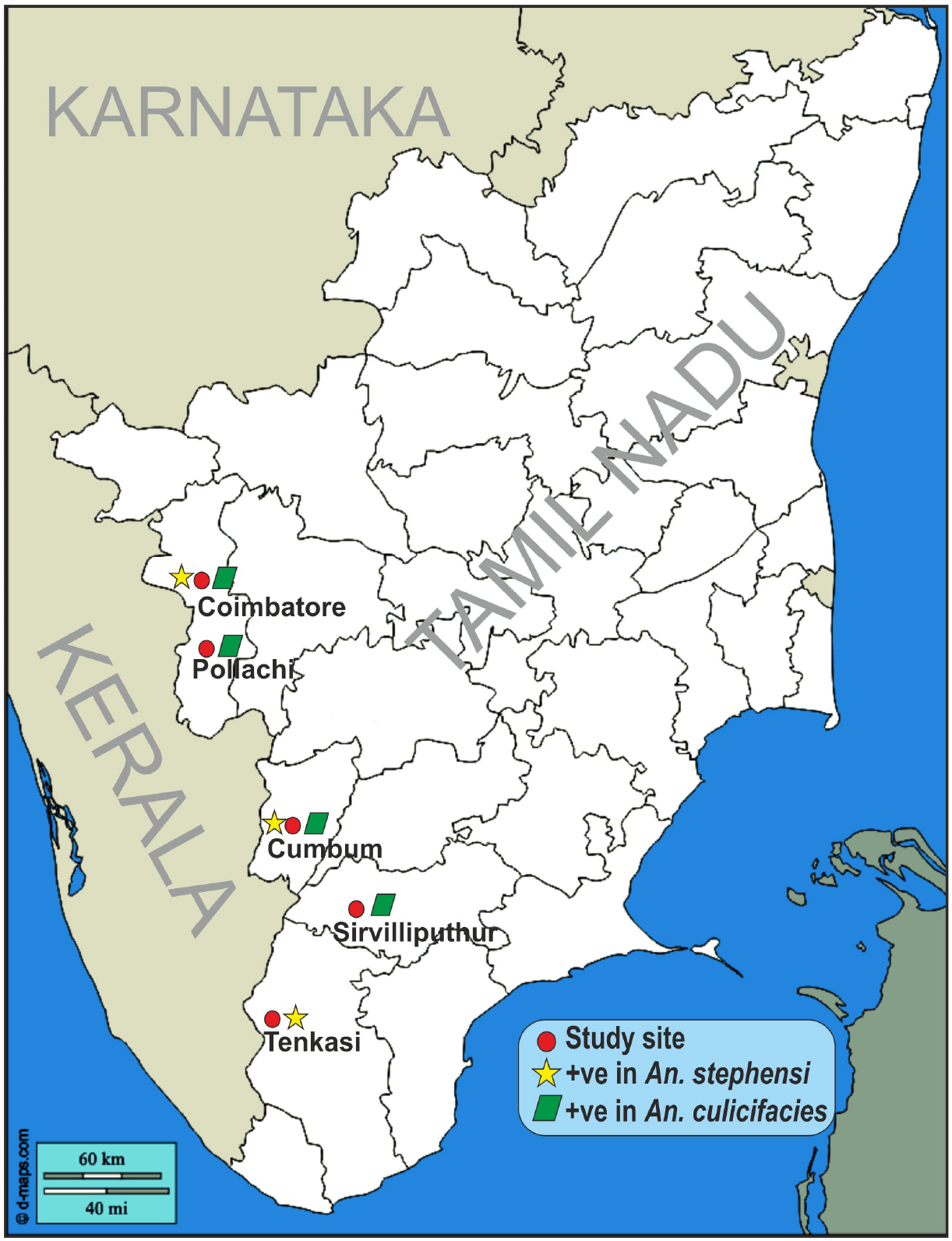
Study site from Tamil Nadu The map shows the study site at 5 different locations along the foothills of the Western Ghats, Southern India. The five locations were Coimbatore, Pollachi, Palani, Cumbum, Sirvilliputhur and Tenkasi. *An. stephensi* and *An. culicifacies* was populated at all the sites (red circle). The presence of *Wolbachia* was marked in yellow star for *An. stephensi* and green diamond for *An. culicifacies*.

### DNA extraction and species identification

DNA isolation from individual mosquito was carried out using the Qiagen Blood and Tissue kit (Qiagen) with slight modification. The initial lysis step post homogenization in PBS was carried-out with proteinase K and lysis buffer at 56°C for 3 hr. Genomic DNA extracted was subjected to *COI* (cytochrome C oxidase subunit I) gene amplification using primers reported by Folmer and colleagues [33] for identification of the mosquito species.

### *Wolbachia* detection by PCR

For *Wolbachia* detection, three different sets of primers targeting conserved genes namely *16S* rRNA gene [34], *Wolbachia* surface protein (*wsp*) gene [35] and *FtsZ* cell cycle gene [36] were used for screening. *Cx. quinquefasciatus* been used as positive control. In addition, for low infection detection, a nested PCR using internal primers targeting 412 bp of *16S* rRNA gene was used [37]. Multilocus strain typing (MLST) of *Wolbachia* was done by targeting five conserved genes *gatB, coxA, hcpA, ftsZ* and *fbpA* as described earlier [38]. The primer details are given in Table S1.

### Molecular phylogenetic studies

The *Wolbachia* positive samples were Sanger sequenced (Barcode Biosciences Pvt. Ltd, Bangalore). The sequencing was carried out from the PCR products of *16S* rRNA (nested PCR) and *COI* from *Wolbachia* and *Anopheles* respectively. The contig assembly was done using MEGA7. The assembled sequences were submitted to GenBank, ENA Database and accession numbers were obtained. Additional mosquito and *Wolbachia* sp. sequences were collected from GenBank to clarify the interspecies relationship. The phylogenetic tree was constructed by maximum likelihood (ML) analysis in MEGA7 software. The tree inference options were set as follows: Heuristic Method Nearest-Neighbor-Interchange (NNI) with the very strong branch swap filter with 1000 bootstrap replicates, gaps were treated as missing. The annotation of phylogenetic tree was performed using iTOL [39]. The number of restriction sites for RsaI was studied using multiple sequence alignment (MSA, Clustal Omega) to find out the supergroup of *Wolbachia* strains reported from this study.

## Results

Mosquito samples were collected from five different study sites (Fig.1). Four hundred and eighty-seven adult female mosquito samples belong to the genus *Anopheles* was screened for the presence of *Wolbachia*. The overall study shows *An. stephensi* (51.75%) population was higher in comparison to *An. culicifacies*. The population of *An. stephensi* was higher in Srivilliputtur (70%), while *An. culicifacies* was higher in Cumbum (69.79%) (Table 1).

**Table 1:** Collection of mosquito samples from different locations

| Place | *An. culicifacies* |  | *An. stephensi* |  |
| --- | --- | --- | --- | --- |
|  | Collected (%*) | *Wolbachia* positive (%<sup>#</sup>) | Collected (%*) | *Wolbachia* positive (%<sup>#</sup>) |
| Coimabtoore | 38 (45.24) | 1 (2.63) | 46 (54.76) | 1 (2.17) |
| Pollachi | 48 (38.71) | 1 (2.08) | 76 (61.29) | 0 |
| Cumbum | 67 (69.79) | 1 (1.49) | 29 (30.21) | 1 (3.44) |
| Srivillputtur | 21 (30.00) | 1 (4.76) | 49 (70.00) | 0 |
| Tenkasi | 61 (53.98) | 0 | 52 (46.02) | 1 (1.92) |
| Total | 235 (48.25) | 4 (1.70) | 252 (51.75) | 3 (1.19) |

\*Percentage from total mosquitoes collected
<sup>#</sup>Percentage of *Wolbachia* positive within species

To identify the presence of *Wolbachia* endosymbiont in mosquito, the genes *wsp* and *FtsZ* genes (Table S1) were amplified; however, no positive results were obtained. MLST by standard primers and protocols did not yield any positive results in all the samples tested (data not shown). Interestingly, nested *16S* PCR amplification targeting the inner region of the *16S* rDNA gene results in positive outcome indicating the presence of *Wolbachia* endosymbiont in *Anopheles* mosquitoes. Out of 487 samples only seven samples i.e., 1.4% were positive for *Wolbachia* endosymbiont, where 3 are from *An. culicifacies* (MN268747, MN268748, MN268749) and 4 from *An. stephensi* (MN268743, MN268744, MN268746 and MN268750) (Table 2). MSA of *Wolbachia* strains reported the presence of four restriction sites (GTAC) for RsaI in agarose gel electrophoresis (data not shown) and *in silico* (Fig.2), indicating the isolates belong to supergroup B.

**Fig.2:**
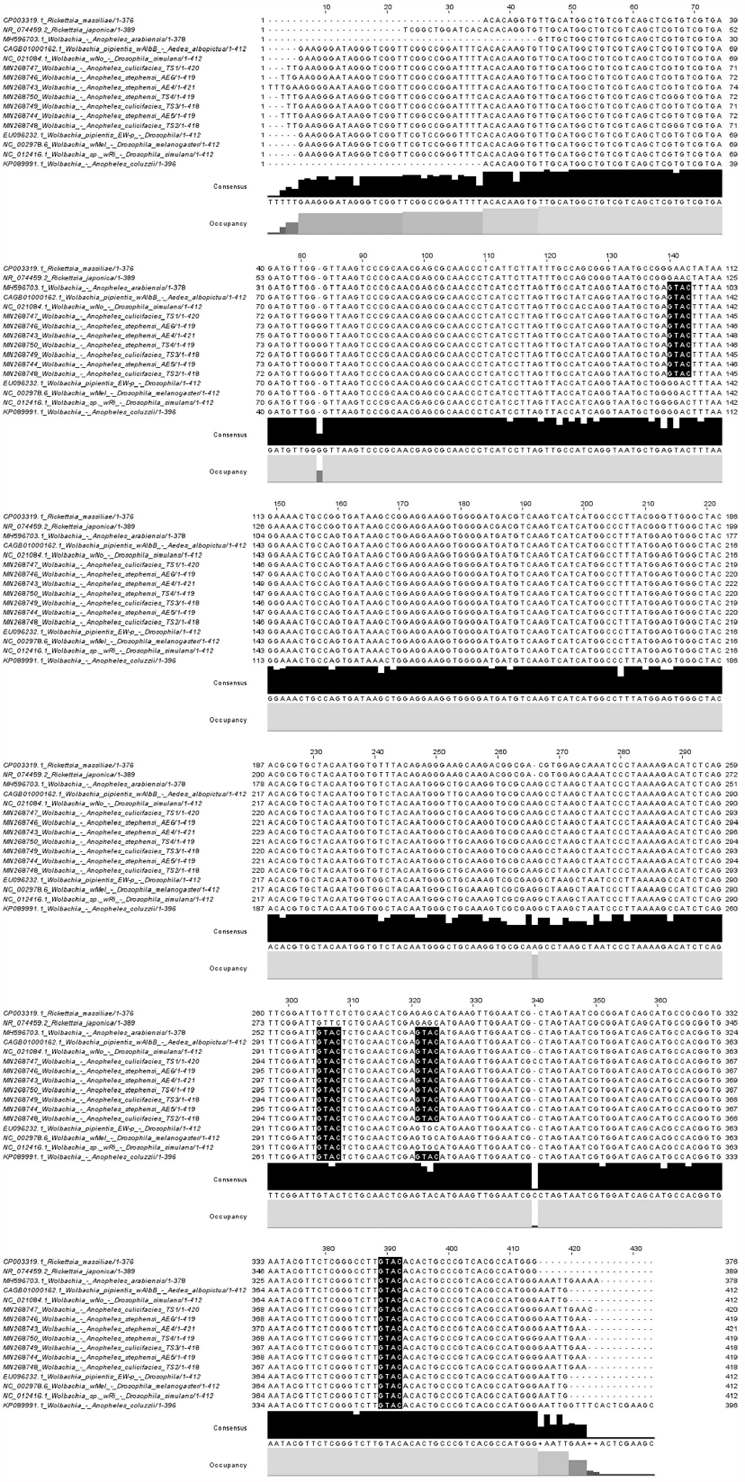
Multiple sequence alignment of *Wolbachia 16S* rRNA sequence The *Wolbachia 16S* rRNA sequence was aligned using Clustal O at EMBL-EBI server and edited in JalView. The restriction site of RsaI was highlighted in black. The sequences used are known *Wolbachia* supergroup A (EU096232, NC_002978.6, NC_012416, KP089991), supergroup B (NC_021984, CAGB01000162, MH967031), outgroup *Ricketssia* (CP003319, NR_074459.2) and our isolates. The restriction sites show our isolates belong to supergroup B with four restriction sites. Supergroup A consists of two restriction sites and, *Ricketssia* consists of a single restriction site.

**Table 2:** The host and the *Wolbachia* endosymbiont isolate ID and sequence accession number

| **Host** | **Isolate ID** | **Mosquito sequence Accession No.** | ***Wolbachia* sequence Accession No.** |
| --- | --- | --- | --- |
| *Anopheles culicifacies* | TS1 | LR736007 | MN268747 |
| *Anopheles culicifacies* | TS2 | LR736008 | MN268748 |
| *Anopheles culicifacies* | TS3 | LR736009 | MN268749 |
| *Anopheles stephensi* | TS4 | LR736010 | MN268750 |
| *Anopheles stephensi* | AE4 | LR736012 | MN268743 |
| *Anopheles stephensi* | AE5 | LR736013 | MN268744 |
| *Anopheles stephensi* | AE6 | LR736014 | MN268746 |

The phylogenetic analysis was done using *Wolbachia* sequences belonging to known Supergroup A, B, C & D. The analysis showed all the *Wolbachia* strains reported from this study (Table S2) were grouped under Supergroup B and non-monophyletic (Fig.3). The *Wolbachia* isolates from this study showed 97.85% - 99.04% sequence similarity with the reference sequences from *Wolbachia* belonging to Supergroup B [CAGB01000162 (wAlbB, France), NC_010981 (wPip, Srilanka), NC021084 (wNo, Sweden), NZ_CP021120 (wMeg, Brazil), NZ_CP034334 (wMau, USA) and NZ_CP034335 (wMau, USA); Table S4]. The genetic diversity within group of Supergroup B (0.2033±0.2139) was higher than supergroup A (0.0012±0.0010) (Table S5). The diversity between supergroup B and other Supergroup A, C & D was highly divergent that ranges from 0.132629±0.1318681 to 0.1225491±0.1292306 (Table S5a).

**Fig.3:**
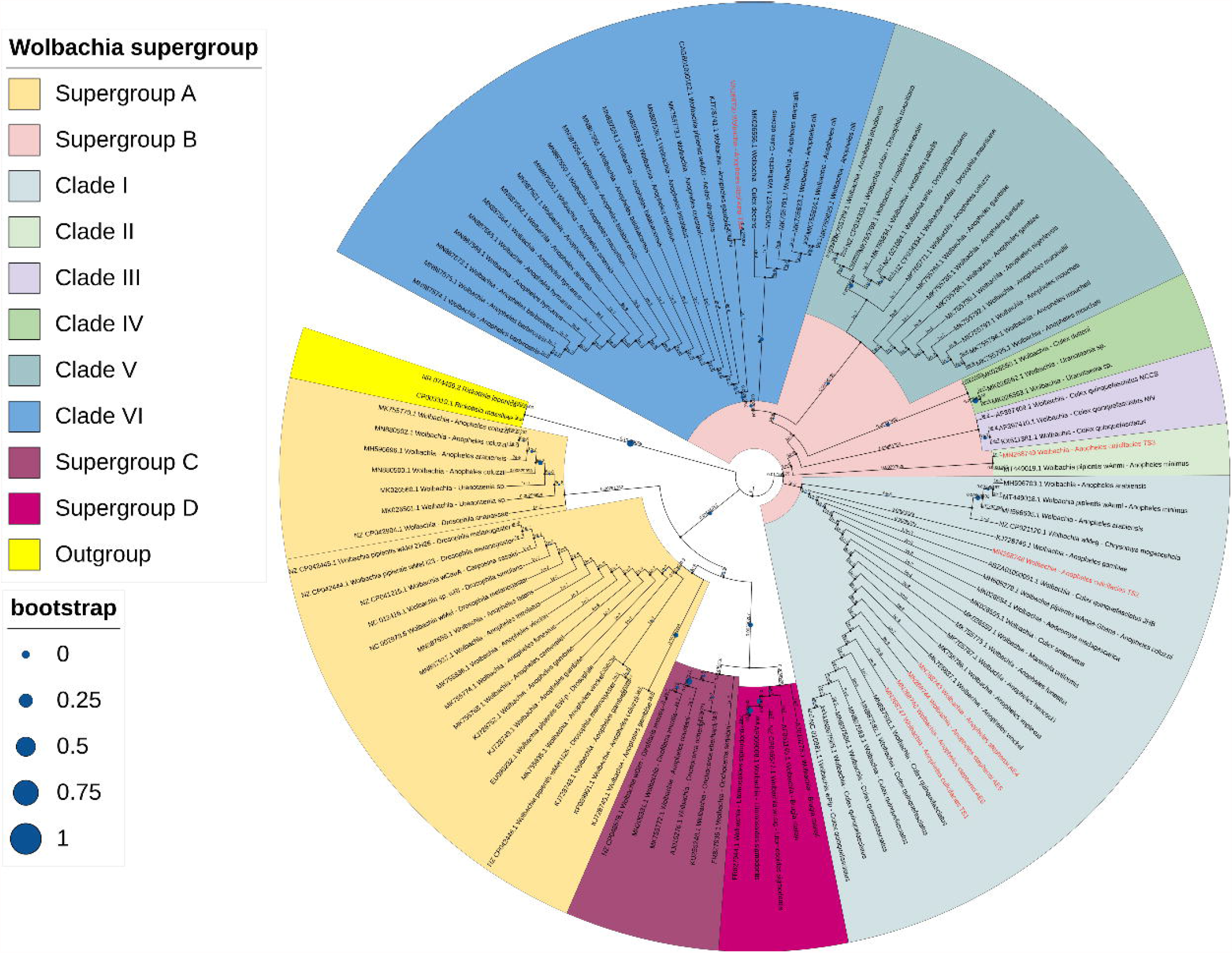
Molecular phylogenetic analysis of *Wolbachia* The phylogeny was inferred from the nucleotide dataset of *16S* rRNA gene by using the ML method. The sequences from this study were represented in red font. The tree with the highest log likelihood (−1044.31) is shown. The analysis involved 102 nucleotide sequences. There were a total of 339 positions in the final dataset. Scale bar 0.02 represents nucleotides substitution per position. The annotation was carried out using iTOL online software.

Based on the divergence, Supergroup B has been subdivided into 6 clades (Fig.3). The isolates from *An. culicifacies* (TS1 & TS2) and *An. stephensi* (AE4, AE5 & AE6), formed a separate clade (Clade I) within supergroup B with the reference strains reported from *Culex quinquefascitus* (NC010981, Srilanka) and *Chrysomya megacephala* (NZ_CP021120, Brazil) (Fig.3). The isolates also shares 98-99% sequence similarity with the two reference sequences (Table S4) and less diverse (0.0018±0.0013; Table S6). The sequence of TS3 is closely related to sequence of *Wolbachia* from *An. minimus* (MT449019; Thailand) with less divergence (0.0014±0.0025) and distinct from other sequence represented as Clade II (Fig.3; Table S6). The sequence of TS4 in Clade VI is closely related with *Wolbachia* sequence reported from *An. gambiae* (KJ728741, Burkina Faso) (Fig.3) with 98.71% sequence similarity and, less diverse (0.0007±0.0010) within Clade VI (Table S6).

All the seven *Wolbachia* positive mosquitoes were amplified using insect *cytochrome C oxidase I* (*COI*) gene (Table S3) and confirmed at species level. The phylogenetic analysis shows there were two clades *An. stephensi* and *An. culicifacies* and, both were supported by 99% bootstrap value (Fig.4). Among the seven *Wolbachia* positive samples, four belongs to *An. stephensi* (LR736010, LR736012, LR736013 and LR736014) and three belongs to *An. culicifacies* s.l. (LR736007 - LR736009) (Table 2). *An. stephensi* showed highest similarity with KX467337, MH538704, KF406680 and NC_028223 (Fig.4) with a high in-group diversity of 0.052±0.01 (Table S7). Interestingly, LR736012 has shown to be divergent from other *An. stephensi* reported. *An. culicifacies* showed high similarity with other *An. culicifacies* strains (KR732656, KJ010898, DQ424962, KJ010896) and supported with 92% bootstrap value (Fig.4) and has low diversity (0.035±0.007) within the analyzed group (Table S7). The diversity between *An. stephensi* and *An. culicifacies* was high (0.16±0.03, Table S8).

**Fig.4:**
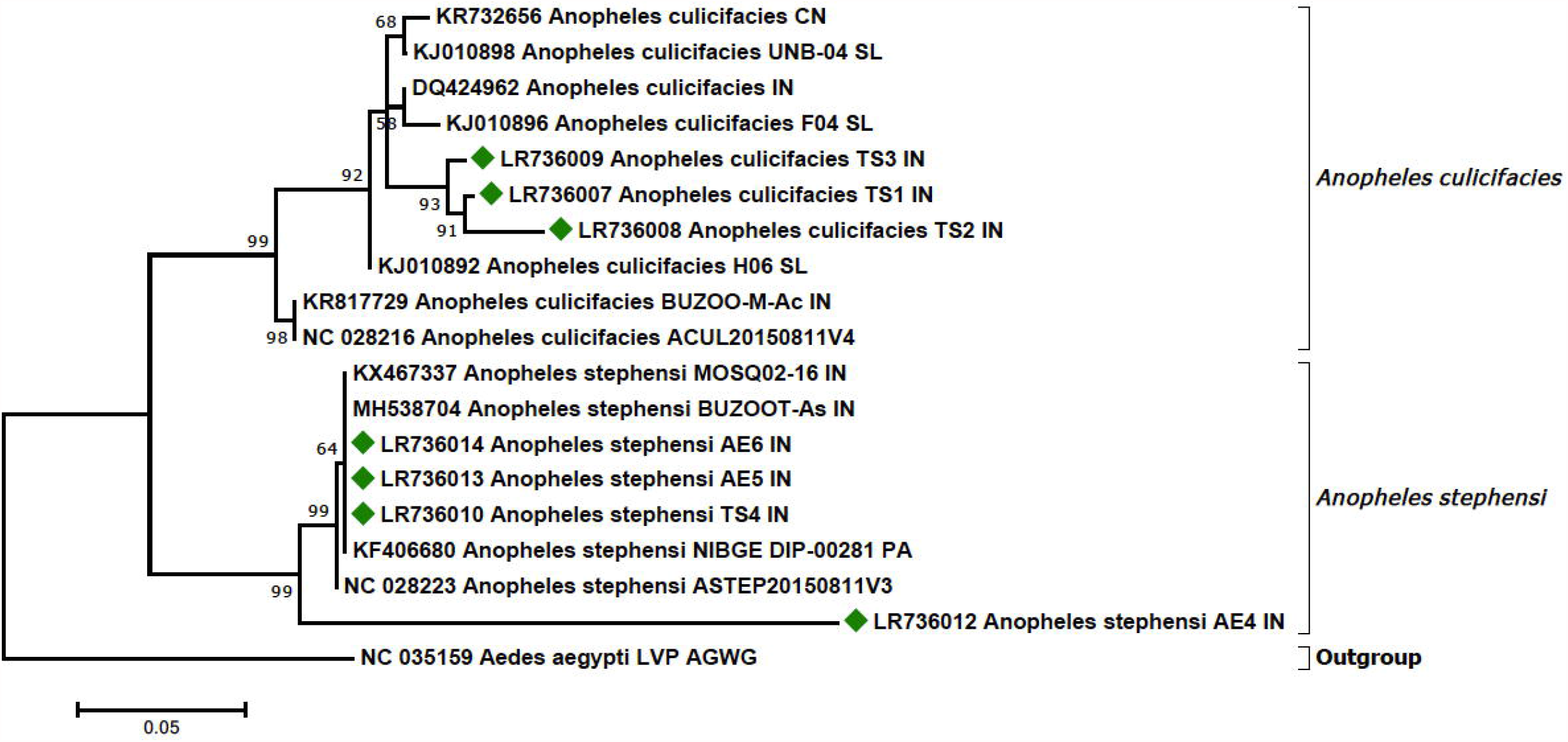
Molecular phylogenetic analysis of *Anopheles* mosquitoes The phylogeny was inferred from the nucleotide dataset of *COI* gene by using the ML method. The sequences from this study were represented as green diamond. The tree with the highest log likelihood (−1429.44) is shown. The analysis involved 19 nucleotide sequences. There were a total of 396 positions in the final dataset. Scale bar 0.05 represents nucleotides substitution per position.

## Discussion

The prevalence of *Wolbachia* has been reported in several arthropods including the orders *Coleoptera, Diptera, Hemiptera, Homoptera, Hymenoptera, Lepidoptera* and *Orthoptera* [5,7]. Among the *Diptera, Wolbachia* is been reported in several *Culicidae* including *Aedes, Culex* and *Coquillettidia* [7,40]. Novel *Wolbachia* infection in *Anopheles* species was least reported [7,40,41]. In this study, we report the occurrence of natural *Wolbachia* endosymbiont in wild *An. stephensi* and *An. culicifacies* for the first time.

### Diversity of isolated *Wolbachia*

*Wolbachia* was identified using *16S* rRNA gene [42,43]; however fine-scale phylogeny was not possible due to low evolutionary divergence of *16S* rRNA, thus *FtsZ* [44] and *wsp* [35] genes were used. Several studies reported amplification with *FtsZ* and *wsp* was unsuccessful [12,45,46] and, our results are similar. Nested PCR amplification of the inner region of *16S* rRNA was used in case of low-intensity *Wolbachia* infection [37] and proved to beneficial in identifying *Wolbachia* infection in *Anopheles* mosquitoes [26,47,37,45]; similarly in our study we detected *Wolbachia* from *An. stephensi* and *An. culicifacies*.

Baldo and colleagues [38] proposed genotyping *Wolbachia* using MLST (*gatB, coxA, hcpA, ftsZ* and *fbpA*). In some cases, MLST in *Wolbachia* were not successful [38] possibly due to primer sequence divergence [13] and low infection densities [27]. Bleidorn and Gerth [48] have pointed out MLST loci are not suitable markers to study either genome-wide divergence rate or strain identification. However in our study, MLST amplification was unsuccessful and, it might be due to low infection density or primer sequence divergence or both.

Werren and colleagues [44] have reported *16S* rRNA can be used to distinguish between A and B supergroup due to the absence of RsaI restriction site in *Wolbachia* belonging to group A. Interestingly, Pourali and colleagues [49] have shown the presence of RsaI restriction site in supergroup A; however they have shown there was more RsaI site in B than A. We have performed *in silico* RsaI restriction site search in *16S* rRNA gene on reported *Wolbachia* supergroup A (EU096232, NC_002978.6, NC_012416, KP089991), supergroup B (NC_021984, CAGB01000162, MH967031) and our isolates; supergroup A has two restriction sites for all strains except KP089991 (*Wolbachia* from *An. coluzzii* [50]) that has three restriction sites, which is possible due to recombination; our isolates and supergroup B shows four restriction sites (Fig.2).

The *16S* rDNA phylogeny shows all the *Wolbachia* isolates from this study belongs to supergroup B and it’s non-monophyletic. Similar to our observation, recent reports [13,51,45] show supergroup B is polyphyletic. The genetic diversity in the clades of supergroup B was less with the excemption of clade III that composed of *Wolbachia* sequence from *Cx. quinquefasciatus* (Fig.3; Table S6). This shows that the nucleotide substitution or recombination was less in *16S* rRNA gene of *Wolbachia* from *Anopheles* mosquitoes.

### Diversity of *Anopheles* mosquitoes and *Wolbachia* prevalence

The genus *Anopheles* belongs to the family *Culicidae* that comprises of 465 species further divided into seven subgenera. *Anopheles* is one among the subgenera consists of 182 species [14,52]. The molecular phylogeny of *Anopheles* was limited to lower level classification respective to malarial vector and, morphologically defined groups found to be monophyletic [52]. Similarly in our study, *An. culicifacies* and *An. stephensi* was identified morphologically; later by *COI* gene amplification and, observed to be monophyletic within their respective species.

To till date, the prevalence of *Wolbachia* has been reported in 20 species of wild *Anopheles* mosquitoes. Baldini and colleagues [12] for the first time reported *Wolbachia* in wild *An. gambiae* and *An. coluzzii* from Burkina Faso. Later *Wolbachia* was reported in other wild *Anopheles* mosquitoes including *An. gambiae* (different from Burkina Faso [13]), *An. arabiensis* [26,27], *An. demeilloni* (previously *An*. species X) [27], *An. moucheti* [27,28], *An. funestus* [47], *An. melas* [53], *An. nili, An. coustani* [28], *An. maculatus* (*s*.*s*.), *An. sawadwongporni, An. pseudowillmori, An. dirus* (*s*.*s*.), *An. baimaii* [51], *An. carnevalei, An. hancocki, An. implexus, An. jebudensis, An. marshallii, An. minimus, An. nigeriensis, An. paludis, An. vinckei* [28], *An. balabacensis, An. latens, An. introlatus, An. macarthuri, An. barbirostris, An. hyrcanus* and *An. sinensis* [45] with our report on additional two species of *Anopheles*, totalling 32 species.

The natural prevalence rate is nil or low in *Anopheles* in comparison to other mosquitoes [41,45,54]. However within *Anopheles* species, the natural *Wolbachia* infection is variable. *An. arabiensis* [26,27], *An. coluzzii* [27,28], *An. funestus* [28,47], *An. melas* [53], *An. maculatus* (*s*.*s*.), *An. sawadwongporni, An. pseudowillmori, An. dirus* (*s*.*s*.), *An. baimaii* [51], *An. balabacensis, An. introlatus, An. macarthuri* [45] shows lesser natural prevalence rate (see Table S9). Higher prevalence rate has been reported from an *An. demeilloni* [27,55,56], *An. moucheti* [27,28], *An. latens, An. hyrcanus* and *An. barbirostris* [45] (see Table S9). *An. gambiae* shows variable prevalence rate [13,27,50] (see Table S9) and, possibly spatial population dynamics [50] may play a role in it. We have lesser prevalence of *Wolbachia* in *An. stephensi* (1.19%) and *An. culicifacies* (1.70%) in comparison to all other reported species.

The less prevalence rate of *Wolbachia* in *Anopheles* mosquitoes has raised several questions. Maternal transmission of *Wolbachia* was observed in natural population of *An. gambiae, An. coluzzii* and *An. arabiensis* [12,13,50,37]. Experimental evidence points out horizontal transfer of *Wolbachia* in *An. gambiae* [57-59], *An. stephensi* [58,60,61] and *An. coluzzii* [37] is possible; however some results in transient infection as in *An. gambiae* [59] than permanent (maternal transmission) as seen in *An. stephensi* [61,60]. Evidence suggests that native microbiome in *Anopheles* mosquitoes impedes the vertical transmission of *Wolbachia* [27,58,62]. *Asaia*, an acetic acid bacterium inhibits *Wolbachia* maternal transmission [58], mutual exclusion of *Wolbachia* in gonads [63] and co-infection of *Asaia* and *Wolbachia* has also been reported [55]. Interestingly, we have also checked for *Asaia* infection using PCR (Table S1) in *Anopheles* mosquitoes including *Wolbachia* positive; but found none (data not shown). *Variovorax*, a beta-proteobacteria has been observed in *Wolbachia* negative *An. coluzzii* [62], which warrants further research as a competitor to *Wolbachia*. The different *Wolbachia* strains can also differ in their interaction with the host [57] probably a reason for lesser prevalence rate. Taken together, prevalence of *Wolbachia* in *Anopheles* mosquito might be subjected to 1) native microbiota interference and 2) *Wolbachia*-host interaction. Since we studied on *Anopheles*, it is possible that native microbiota could interfere in *Wolbachia* colonizing this species leading to lesser prevalence rate; further research will be carried out in the future.

Lateral gene transfer (LGT) of *Wolbachia* genome was observed in *Callosobruchus chinensis* [64,65], *Onchocerca volvulus* [66] and, *D. ananassae* [67]. Incase of mosquitoes, LGT has been observed in *Ae. aegypti, Ae. mascarensis* [68] and *An. gambiae* [69]. Salivary gland surface (*SGS*) genes from *An. gambiae* and *Ae. aegypti* genome is said to be transferred from *Wolbachia* via LGT [69], where these genes are particularly found in female salivary gland; it’s expression increase with age, after blood feeding and, facilitates *Plasmodium* sporozoite invasion [70,69]. However, there is no study on SGS relationship with *Wolbachia* infection. Endogenous bornavirus-like nucleoprotein, a functional protein homologous to Borna virus *nucleoprotein* integrated into *Ictidomys tridecemlineatus* ∼8.5 MYA [71] inhibits exogenous Borna virus *in vivo* [72]. Similarly we hypothesis, *SGS* gene might interfere with *Wolbachia* infection in *Anopheles* mosquitoes leads to less prevalence rate which needs further research.

### Is *Wolbachia* a contaminant in *Anopheles* mosquitoes?

According to Chrostek and Gerth [73], *Wolbachia* in *An. gambiae* could be from contamination through several sources – 1) *Wolbachia* present in ectoparasitic mites or midges and endoparasitic nematodes, 2) *Wolbachia* contamination from plants, where *Wolbachia* infested insects feed on the plants might transfer to uninfected insects reared on the same plants and 3) cohabitating insect with *Wolbachia* contaminating the water bodies, thus infecting other mosquito species. However, recent report evidently shows *Wolbachia* as an endosymbiont in *Anopheles* species – *An. gambiae, An. melas* [55], *An. moucheti* and *An. demeilloni* [56].

In our case, we would like to clarify the *Wolbachia* is from *Anopheles* mosquitoes and not from other sources. 1) Only adult female *An. culicifacies* and *An. stephansi* with no ectoparasitic mites or midges were collected from the study sites. In case of contaminant from filarial nematode, the *Wolbachia* should be from other Supergroup such as C and D. According to Ayala and colleagues [28], *Wolbachia* strains from *An. coustani* belongs to Supergroup B & C; further investigation shows *Wolbachia* belonging to Supergroup C (MK755772) is from nematode *Dirofilaria immitis* infected *An. coustani*. Our observation on the sequence (MK755772) shows the same that it belongs to Supergroup C with a 94% bootstrap (see Fig.3). This indicates *Wolbachia* from filarial parasites is distinct and belong to Supergroup C/D [74-76] and can be used as an indicator of *Wolbachia* from filarial endoparasites. In our study, *Wolbachia* are from Supergroup B and none from Supergroup C/D (see Fig.3), indicating all the *Wolbachia* strains are from *Anopheles* mosquitoes than from filarial nematodes. Thus it eliminates the possibility of *Wolbachia* from ecto- and endo-parasites. 2) Adult female *An. culicifacies* and *An. stephensi* usually feed on blood meal [77,78]; the collected mosquitoes were surface sterilized and preserved in 95% ethanol and, no laboratory breeding was performed indicating *Wolbachia* is not from infected plants or other environmental contaminants. 3) The majority of co-habitating mosquito with *An. stephensi* and *An. culicifacies* is *Cx. quinquefaciatus* (personal communication). Few studies from India reported *Wolbachia* in *Cx. quinquefaciatus* [79-81]. We have included *Cx. quinquefaciatus* as positive control in our PCR experiments and were positive for *Wolbachia* (data not shown). When the *Wolbachia* sequences of *Cx. quequefasciatus* (AF397409, AF397410) from India was phylogenetically analyzed it forms a distinctive separate clade and not closer to our isolates (Fig.3). Even if *Wolbachia* is from *Cx. quinquefasciatus*, it can only consider as infection rather than ‘contamination’; the infection might happened at the larval stages and the co-existence in the adult stage (our isolates are from adult female mosquitoes) should be consider as symbiotic relationship. In future, studies on cohabitation and *Wolbachia* transmission will be carried. Based on the above facts, we confirm *Wolbachia* in our study is from *An. stephensi* and *An. culicifacies* and not from ecto- or endo-parasites or other sources.

Chrostek and Gerth [73] pointed out that true symbiosis has to be established by demonstrating intercellular bacterial cells and intraovarian transmission. To till date, the available techniques are limited and thus addressing the above said factors in ‘wild mosquitoes’ is quite a challenge, but the future might hold a better way to prove the above said factors in elucidating true symbiosis of *Wolbachia* in wild mosquitoes.

## Conclusion

The current study has shown *Wolbachia* for the first time in *Anopheles* mosquitoes namely *An. culicifacies* and *An. stephensi* from Tamil Nadu, India. Nested *16S* rRNA PCR amplification is helpful in identification than *wsp, FtsZ* and MLST loci genes. The prevalence is lesser compared to other mosquitoes, which may be due to inhibition by native microbiota, host and *Wolbachia* interaction, and/or inhibition by endogenous gene product that result from LGT; these factors will be analyzed in our future research. Future studies will also include the different organs of the mosquito showing *Wolbachia* by FISH method.

## Acknowledgements

SGS would like to acknowledge DST-SERB (YSS/2015/001847) and DHR-HRD (DHR/HRD/YS-14-2015-16) fellowships. The authors acknowledge DBT-BIF Centre, Lady Doak College for providing bioinformatics facility for data analysis.

## Declarations

## Author contribution

Concept, study design and methodology: SGS and APA. Performed lab experiments: TM and APA. Phylogenetic design and analysis: TM, SGS and APA. Pre-draft written: TM, SGS and APA. Critical revision of final version: SGS and APA. All authors read and approved the final manuscript.

## Availability of data and materials

The data are provided in the supplementary section and the sequence data from this study is available in GenBank.

## Reference

1. Hertig M, Wolbach SB (1924) Studies on Rickettsia-Like Micro-Organisms in Insects. The Journal of medical research 44 (3):329–374

2. Werren JH (1997) Biology of Wolbachia. Annual review of entomology 42:587–609. doi:10.1146/annurev.ento.42.1.587

3. Werren JH, Baldo L, Clark ME (2008) Wolbachia: master manipulators of invertebrate biology. Nature reviews Microbiology 6 (10):741–751. doi:10.1038/nrmicro1969

4. Hilgenboecker K, Hammerstein P, Schlattmann P, Telschow A, Werren JH (2008) How many species are infected with Wolbachia?--A statistical analysis of current data. FEMS microbiology letters 281 (2):215–220. doi:10.1111/j.1574-6968.2008.01110.x

5. Werren JH, Windsor D, Guo LR (1995) Distribution of Wolbachia among neotropical arthropods. Proc Royal Soc Lond B 262 (1364):197–204. doi:doi:10.1098/rspb.1995.0196

6. Sicard M, Bonneau M, Weill M (2019) Wolbachia prevalence, diversity, and ability to induce cytoplasmic incompatibility in mosquitoes. Current opinion in insect science 34:12–20. doi:10.1016/j.cois.2019.02.005

7. de Oliveira CD, Goncalves DS, Baton LA, Shimabukuro PH, Carvalho FD, Moreira LA (2015) Broader prevalence of Wolbachia in insects including potential human disease vectors. Bulletin of entomological research 105 (3):305–315. doi:10.1017/S0007485315000085

8. Mahilum MM, Storch V, Becker N (2003) Molecular and electron microscopic identification of Wolbachia in Culex pipiens complex populations from the Upper Rhine Valley, Germany, and Cebu City, Philippines. Journal of the American Mosquito Control Association 19 (3):206–210

9. Dutton TJ, Sinkins SP (2004) Strain-specific quantification of Wolbachia density in Aedes albopictus and effects of larval rearing conditions. Insect molecular biology 13 (3):317–322. doi:10.1111/j.0962-1075.2004.00490.x

10. Sinkins SP, Braig HR, O’Neill SL (1995) Wolbachia pipientis: bacterial density and unidirectional cytoplasmic incompatibility between infected populations of Aedes albopictus. Experimental parasitology 81 (3):284–291. doi:10.1006/expr.1995.1119

11. Coon KL, Brown MR, Strand MR (2016) Mosquitoes host communities of bacteria that are essential for development but vary greatly between local habitats. Molecular ecology 25 (22):5806–5826. doi:10.1111/mec.13877

12. Baldini F, Segata N, Pompon J, Marcenac P, Shaw WR, Dabire RK, Diabate A, Levashina EA, Catteruccia F (2014) Evidence of natural Wolbachia infections in field populations of Anopheles gambiae. Nature communications 5:3985. doi:10.1038/ncomms4985

13. Gomes FM, Hixson BL, Tyner MDW, Ramirez JL, Canepa GE, Alves ESTL, Molina-Cruz A, Keita M, Kane F, Traore B, Sogoba N, Barillas-Mury C (2017) Effect of naturally occurring Wolbachia in Anopheles gambiae s.l. mosquitoes from Mali on Plasmodium falciparum malaria transmission. Proceedings of the National Academy of Sciences of the United States of America 114 (47):12566–12571. doi:10.1073/pnas.1716181114

14. Harbach RE (2013) The Phylogeny and Classification of Anopheles. In: Manguin S (ed) Anopheles mosquitoes - New insights into malaria vectors. IntechOpen. doi:10.5772/54695

15. Hay SI, Sinka ME, Okara RM, Kabaria CW, Mbithi PM, Tago CC, Benz D, Gething PW, Howes RE, Patil AP, Temperley WH, Bangs MJ, Chareonviriyaphap T, Elyazar IR, Harbach RE, Hemingway J, Manguin S, Mbogo CM, Rubio-Palis Y, Godfray HC (2010) Developing global maps of the dominant anopheles vectors of human malaria. PLoS medicine 7 (2):e1000209. doi:10.1371/journal.pmed.1000209

16. Sinka ME, Bangs MJ, Manguin S, Chareonviriyaphap T, Patil AP, Temperley WH, Gething PW, Elyazar IR, Kabaria CW, Harbach RE, Hay SI (2011) The dominant Anopheles vectors of human malaria in the Asia-Pacific region: occurrence data, distribution maps and bionomic precis. Parasites & vectors 4:89. doi:10.1186/1756-3305-4-89

17. WHO (2019) World malaria report 2019. World Health Organization, Geneva

18. Subbarao SK, Nanda N, Rahi M, Raghavendra K (2019) Biology and bionomics of malaria vectors in India: existing information and what more needs to be known for strategizing elimination of malaria. Malaria journal 18 (1):396. doi:10.1186/s12936-019-3011-8

19. Dev V, Sharma VP (2013) The dominant mosquito vectors of human malaria in India. In: Manguin S (ed) Anopheles mosquitoes - New insights into malaria vectors. IntechOpen. doi:10.5772/55215

20. Goswami G, Singh OP, Nanda N, Raghavendra K, Gakhar SK, Subbarao SK (2006) Identification of all members of the anopheles culicifacies complex using allele-specific polymerase chain reaction assays. The American journal of tropical medicine and hygiene 75 (3):454–460

21. Sharma SK, Hamzakoya KK (2001) Geographical Spread of Anopheles stephensi Vector of Urban Malaria, and Aedes aegypti, Vector of Dengue/DHF, in the Arabian Sea Islands of Lakshadweep, India. Dengue Bulletin 25:88–91

22. Arjunan NK, Kadarkarai M, Kumar S, Pari M, Thiyagarajan N, Vincent CT, Barnard DR (2015) Factors influencing the spatial distribution of Anopheles larvae in Coimbatore District, Tamil Nadu, India. Acta tropica 152:121–130. doi:10.1016/j.actatropica.2015.09.007

23. Surendran SN, Sivabalakrishnan K, Sivasingham A, Jayadas TTP, Karvannan K, Santhirasegaram S, Gajapathy K, Senthilnanthanan M, Karunaratne SP, Ramasamy R (2019) Anthropogenic factors driving recent range expansion of the malaria vector Anopheles stephensi. Frontiers in public health 7:53. doi:10.3389/fpubh.2019.00053

24. Suguna SG, Tewari SC, Mani TR, Hiriyan J, Reuben R (1983) Anopheles culicifacies species complex in Thenpennaiyar riverine tract, Tamil Nadu. The Indian journal of medical research 77:455–459

25. Kar I, Subbarao SK, Eapen A, Ravindran J, Satyanarayana TS, Raghavendra K, Nanda N, Sharma VP (1999) Evidence for a new malaria vector species, species E, within the Anopheles culicifacies complex (Diptera: Culicidae). Journal of medical entomology 36 (5):595–600. doi:10.1093/jmedent/36.5.595

26. Baldini F, Rouge J, Kreppel K, Mkandawile G, Mapua SA, Sikulu-Lord M, Ferguson HM, Govella N, Okumu FO (2018) First report of natural Wolbachia infection in the malaria mosquito Anopheles arabiensis in Tanzania. Parasites & vectors 11 (1):635. doi:10.1186/s13071-018-3249-y

27. Jeffries CL, Lawrence GG, Golovko G, Kristan M, Orsborne J, Spence K, Hurn E, Bandibabone J, Tantely LM, Raharimalala FN, Keita K, Camara D, Barry Y, Wat’senga F, Manzambi EZ, Afrane YA, Mohammed AR, Abeku TA, Hedge S, Khanipov K, Pimenova M, Fofanov Y, Boyer S, Irish SR, Hughes GL, Walker T (2018) Novel Wolbachia strains in Anopheles malaria vectors from Sub-Saharan Africa. Wellcome open research 3:113. doi:10.12688/wellcomeopenres.14765.2

28. Ayala D, Akone-Ella O, Rahola N, Kengne P, Ngangue MF, Mezeme F, Makanga BK, Nigg M, Costantini C, Simard F, Prugnolle F, Roche B, Duron O, Paupy C (2019) Natural Wolbachia infections are common in the major malaria vectors in Central Africa. Evolutionary Applications 12 (8):1583–1594. doi:10.1111/eva.12804

29. Christophers SR (1933) The fauna of British India, including Ceylon and Burma. Diptera.Vol. IV. Family Culicidae. Tribe Anophelini. Taylor and Francis, London

30. Das BP, Rajagopal R, Akiyama J (1990) Pictorial key to the species of Indian anopheline mosquitoes. Pictorial key to the species of Indian anopheline mosquitoes. Journal of Pure and Applied Zoology 2 (3):131–162

31. Pennington MJ, Prager SM, Walton WE, Trumble JT (2016) Culex quinquefasciatus larval microbiomes vary with instar and exposure to common wastewater contaminants. Scientific Reports 6 (1):21969. doi:10.1038/srep21969

32. Reeves LE, Holderman CJ, Gillett-Kaufman JL, Kawahara AY, Kaufman PE (2016) Maintenance of host DNA integrity in field-preserved mosquito (Diptera: Culicidae) blood meals for identification by DNA barcoding. Parasites & vectors 9 (1):503. doi:10.1186/s13071-016-1791-z

33. Folmer O, Black M, Hoeh W, Lutz R, Vrijenhoek R (1994) DNA primers for amplification of mitochondrial cytochrome c oxidase subunit I from diverse metazoan invertebrates. Molecular marine biology and biotechnology 3 (5):294–299

34. Werren JH, Windsor DM (2000) Wolbachia infection frequencies in insects: evidence of a global equilibrium? Proceedings Biological sciences 267 (1450):1277–1285. doi:10.1098/rspb.2000.1139

35. Zhou W, Rousset F, O’Neil S (1998) Phylogeny and PCR-based classification of Wolbachia strains using wsp gene sequences. Proceedings Biological sciences 265 (1395):509–515. doi:10.1098/rspb.1998.0324

36. Werren JH, Jaenike J (1995) Wolbachia and cytoplasmic incompatibility in mycophagous Drosophila and their relatives. Heredity 75 (Pt 3):320–326. doi:10.1038/hdy.1995.140

37. Shaw WR, Marcenac P, Childs LM, Buckee CO, Baldini F, Sawadogo SP, Dabire RK, Diabate A, Catteruccia F (2016) Wolbachia infections in natural Anopheles populations affect egg laying and negatively correlate with Plasmodium development. Nature communications 7:11772. doi:10.1038/ncomms11772

38. Baldo L, Dunning Hotopp JC, Jolley KA, Bordenstein SR, Biber SA, Choudhury RR, Hayashi C, Maiden MC, Tettelin H, Werren JH (2006) Multilocus sequence typing system for the endosymbiont Wolbachia pipientis. Applied and environmental microbiology 72 (11):7098–7110. doi:10.1128/AEM.00731-06

39. Letunic I, Bork P (2021) Interactive Tree Of Life (iTOL) v5: an online tool for phylogenetic tree display and annotation. Nucleic Acids Research. doi:10.1093/nar/gkab301

40. Ricci I, Cancrini G, Gabrielli S, D’Amelio S, Favi G (2002) Searching for Wolbachia (Rickettsiales: Rickettsiaceae) in mosquitoes (Diptera: Culicidae): large polymerase chain reaction survey and new identifications. Journal of medical entomology 39 (4):562–567. doi:10.1603/0022-2585-39.4.562

41. Kittayapong P, Baisley KJ, Baimai V, O’Neill SL (2000) Distribution and diversity of Wolbachia infections in Southeast Asian mosquitoes (Diptera: Culicidae). Journal of medical entomology 37 (3):340–345. doi:10.1093/jmedent/37.3.340

42. O’Neill SL, Giordano R, Colbert AM, Karr TL, Robertson HM (1992) 16S rRNA phylogenetic analysis of the bacterial endosymbionts associated with cytoplasmic incompatibility in insects. Proceedings of the National Academy of Sciences of the United States of America 89 (7):2699–2702. doi:10.1073/pnas.89.7.2699

43. Rousset F, Vautrin D, Solignac M (1992) Molecular identification of Wolbachia, the agent of cytoplasmic incompatibility in Drosophila simulans, and variability in relation with host mitochondrial types. Proceedings Biological sciences 247 (1320):163–168. doi:10.1098/rspb.1992.0023

44. Werren JH, Zhang W, Guo LR (1995) Evolution and phylogeny of Wolbachia: reproductive parasites of arthropods. Proceedings Biological sciences 261 (1360):55–63. doi:10.1098/rspb.1995.0117

45. Wong ML, Liew JWK, Wong WK, Pramasivan S, Mohamed Hassan N, Wan Sulaiman WY, Jeyaprakasam NK, Leong CS, Low VL, Vythilingam I (2020) Natural Wolbachia infection in field-collected Anopheles and other mosquito species from Malaysia. Parasites & vectors 13 (1):414. doi:10.1186/s13071-020-04277-x

46. Marcon HS, Coscrato VE, Selivon D, Perondini AL, Marino CL (2011) Variations in the sensitivity of different primers for detecting Wolbachia in Anastrepha (diptera: tephritidae). Brazilian journal of microbiology : [publication of the Brazilian Society for Microbiology] 42 (2):778–785. doi:10.1590/S1517-838220110002000046

47. Niang EHA, Bassene H, Makoundou P, Fenollar F, Weill M, Mediannikov O (2018) First report of natural Wolbachia infection in wild Anopheles funestus population in Senegal. Malaria journal 17 (1):408. doi:10.1186/s12936-018-2559-z

48. Bleidorn C, Gerth M (2018) A critical re-evaluation of multilocus sequence typing (MLST) efforts in Wolbachia. FEMS Microbiol Ecol 94 (1). doi:10.1093/femsec/fix163

49. Pourali P, Roayaei Ardakani M, Jolodar A, Razi Jalali MH (2009) PCR screening of the Wolbachia in some arthropods and nematodes in Khuzestan province. Iranian Journal of Veterinary Research 10 (3):216–222

50. Buck M, Nilsson LK, Brunius C, Dabiré RK, Hopkins R, Terenius O (2016) Bacterial associations reveal spatial population dynamics in Anopheles gambiae mosquitoes. Sci Rep 6:22806. doi:10.1038/srep22806

51. Sawasdichai S, Chaumeau V, Dah T, Kulabkeeree T, Kajeechiwa L, Phanaphadungtham M, Trakoolchengkaew M, Kittiphanakun P, Akararungrot Y, Oo K, Delmas G, White NJ, Nosten FH (2019) Detection of diverse Wolbachia 16S rRNA sequences at low titers from malaria vectors in Kayin state, Myanmar. Wellcome open research 4:11. doi:10.12688/wellcomeopenres.15005.4

52. Harbach RE (2004) The classification of genus Anopheles (Diptera: Culicidae): a working hypothesis of phylogenetic relationships. Bulletin of entomological research 94 (6):537–553. doi:10.1079/ber2004321

53. Jeffries CL, Cansado-Utrilla C, Stica C, Walker T (2019) High density Novel Wolbachia strains in Anopheles species from Guinea. bioRxiv:772855. doi:10.1101/772855

54. Rasgon JL, Scott TW (2004) An initial survey for Wolbachia (Rickettsiales: Rickettsiaceae) infections in selected California mosquitoes (Diptera: Culicidae). Journal of medical entomology 41 (2):255–257. doi:10.1603/0022-2585-41.2.255

55. Jeffries CL, Cansado-Utrilla C, Beavogui AH, Stica C, Lama EK, Kristan M, Irish SR, Walker T (2020) Evidence for natural hybridisation and novel Wolbachia strain superinfections in the Anopheles gambiae complex from Guinea. bioRxiv:772855. doi:10.1101/772855

56. Walker T, Quek S, Jeffries CL, Bandibabone J, Dhokiya V, Bamou R, Kristan M, Messenger LA, Gidley A, Hornett EA, Anderson ER, Cansado-Utrilla C, Hegde S, Bantuzeko C, Stevenson JC, Lobo NF, Wagstaff SC, Antonio Nkondjio C, Heinz E, Hughes GL (2020) Genomic and microscopic evidence of stable high density and maternally inherited Wolbachia infections in Anopheles mosquitoes. bioRxiv:2020.2010.2029.357400. doi:10.1101/2020.10.29.357400

57. Hughes GL, Vega-Rodriguez J, Xue P, Rasgon JL (2012) Wolbachia strain wAlbB enhances infection by the rodent malaria parasite Plasmodium berghei in Anopheles gambiae mosquitoes. Applied and environmental microbiology 78 (5):1491–1495. doi:10.1128/AEM.06751-11

58. Hughes GL, Dodson BL, Johnson RM, Murdock CC, Tsujimoto H, Suzuki Y, Patt AA, Cui L, Nossa CW, Barry RM, Sakamoto JM, Hornett EA, Rasgon JL (2014) Native microbiome impedes vertical transmission of Wolbachia in Anopheles mosquitoes. Proceedings of the National Academy of Sciences of the United States of America 111 (34):12498–12503. doi:10.1073/pnas.1408888111

59. Hughes GL, Koga R, Xue P, Fukatsu T, Rasgon JL (2011) Wolbachia infections are virulent and inhibit the human malaria parasite Plasmodium falciparum in Anopheles gambiae. PLoS pathogens 7 (5):e1002043. doi:10.1371/journal.ppat.1002043

60. Bian G, Joshi D, Dong Y, Lu P, Zhou G, Pan X, Xu Y, Dimopoulos G, Xi Z (2013) Wolbachia invades Anopheles stephensi populations and induces refractoriness to Plasmodium infection. Science 340 (6133):748–751. doi:10.1126/science.1236192

61. Joshi D, Pan X, McFadden MJ, Bevins D, Liang X, Lu P, Thiem S, Xi Z (2017) The maternally inheritable Wolbachia wAlbB induces refractoriness to Plasmodium berghei in Anopheles stephensi. Frontiers in microbiology 8 (366). doi:10.3389/fmicb.2017.00366

62. Straub TJ, Shaw WR, Marcenac P, Sawadogo SP, Dabire RK, Diabate A, Catteruccia F, Neafsey DE (2020) The Anopheles coluzzii microbiome and its interaction with the intracellular parasite Wolbachia. Sci Rep 10 (1):13847. doi:10.1038/s41598-020-70745-0

63. Rossi P, Ricci I, Cappelli A, Damiani C, Ulissi U, Mancini MV, Valzano M, Capone A, Epis S, Crotti E, Chouaia B, Scuppa P, Joshi D, Xi Z, Mandrioli M, Sacchi L, O’Neill SL, Favia G (2015) Mutual exclusion of Asaia and Wolbachia in the reproductive organs of mosquito vectors. Parasites & vectors 8 (1):278. doi:10.1186/s13071-015-0888-0

64. Kondo N, Nikoh N, Ijichi N, Shimada M, Fukatsu T (2002) Genome fragment of Wolbachia endosymbiont transferred to X chromosome of host insect. Proceedings of the National Academy of Sciences of the United States of America 99 (22):14280–14285. doi:10.1073/pnas.222228199

65. Nikoh N, Tanaka K, Shibata F, Kondo N, Hizume M, Shimada M, Fukatsu T (2008) Wolbachia genome integrated in an insect chromosome: Evolution and fate of laterally transferred endosymbiont genes. Genome Research 18 (2):272–280. doi:10.1101/gr.7144908

66. Fenn K, Conlon C, Jones M, Quail MA, Holroyd NE, Parkhill J, Blaxter M (2006) Phylogenetic Relationships of the Wolbachia of Nematodes and Arthropods. PLoS pathogens 2 (10):e94. doi:10.1371/journal.ppat.0020094

67. Hotopp JCD, Clark ME, Oliveira DCSG, Foster JM, Fischer P, Torres MCM, Giebel JD, Kumar N, Ishmael N, Wang S, Ingram J, Nene RV, Shepard J, Tomkins J, Richards S, Spiro DJ, Ghedin E, Slatko BE, Tettelin H, Werren JH (2007) Widespread Lateral Gene Transfer from Intracellular Bacteria to Multicellular Eukaryotes. Science 317 (5845):1753–1756. doi:10.1126/science.1142490

68. Klasson L, Kambris Z, Cook PE, Walker T, Sinkins SP (2009) Horizontal gene transfer between Wolbachia and the mosquito Aedes aegypti. BMC Genomics 10 (1):33. doi:10.1186/1471-2164-10-33

69. Korochkina S, Barreau C, Pradel G, Jeffery E, Li J, Natarajan R, Shabanowitz J, Hunt D, Frevert U, Vernick KD (2006) A mosquito-specific protein family includes candidate receptors for malaria sporozoite invasion of salivary glands. Cellular Microbiology 8 (1):163–175. doi:10.1111/j.1462-5822.2005.00611.x

70. King JG, Vernick KD, Hillyer JF (2011) Members of the salivary gland surface protein (SGS) family are major immunogenic components of mosquito saliva. The Journal of biological chemistry 286 (47):40824–40834. doi:10.1074/jbc.M111.280552

71. Suzuki Y, Kobayashi Y, Horie M, Tomonaga K (2014) Origin of an endogenous bornavirus-like nucleoprotein element in thirteen-lined ground squirrels. Genes & genetic systems 89 (3):143–148. doi:10.1266/ggs.89.143

72. Fujino K, Horie M, Honda T, Merriman DK, Tomonaga K (2014) Inhibition of Borna disease virus replication by an endogenous bornavirus-like element in the ground squirrel genome. Proceedings of the National Academy of Sciences 111 (36):13175–13180. doi:10.1073/pnas.1407046111

73. Chrostek E, Gerth M (2019) Is Anopheles gambiae a Natural Host of Wolbachia? mBio 10 (3). doi:10.1128/mBio.00784-19

74. Inácio da Silva LM, Dezordi FZ, Paiva MHS, Wallau GL (2021) Systematic Review of Wolbachia Symbiont Detection in Mosquitoes: An Entangled Topic about Methodological Power and True Symbiosis. Pathogens 10 (1):39

75. Ros VID, Fleming VM, Feil EJ, Breeuwer JAJ (2009) How Diverse Is the Genus Wolbachia? Multiple-Gene Sequencing Reveals a Putatively New Wolbachia Supergroup Recovered from Spider Mites (Acari: Tetranychidae). Applied and environmental microbiology 75 (4):1036–1043. doi:10.1128/aem.01109-08

76. Bouchery T, Lefoulon E, Karadjian G, Nieguitsila A, Martin C (2013) The symbiotic role of Wolbachia in Onchocercidae and its impact on filariasis. Clinical microbiology and infection : the official publication of the European Society of Clinical Microbiology and Infectious Diseases 19 (2):131–140. doi:10.1111/1469-0691.12069

77. Swami KK, Srivastava M (2012) Blood Meal Preference of Some Anopheline Mosquitoes in Command and Non-command Areas of Rajasthan, India. Journal of arthropod-borne diseases 6 (2):98–103

78. Thomas S, Ravishankaran S, Justin NAJA, Asokan A, Mathai MT, Valecha N, Montgomery J, Thomas MB, Eapen A (2017) Resting and feeding preferences of Anopheles stephensi in an urban setting, perennial for malaria. Malaria journal 16 (1):111. doi:10.1186/s12936-017-1764-5

79. Sunish IP, Rajendran R, Paramasivan R, Dhananjeyan KJ, Tyagi BK (2011) Wolbachia endobacteria in a natural population of Culex quinquefasciatus from filariasis endemic villages of south India and its phylogenetic implication. Trop Biomed 28 (3):569–576

80. Muniaraj M, Paramasivan R, Sunish IP, Arunachalam N, Mariappan T, Jerald Leo SV, Dhananjeyan KJ (2012) Detection of Wolbachia endobacteria in Culex quinquefasciatus by Gimenez staining and confirmation by PCR. Journal of vector borne diseases 49 (4):258–261

81. Pidiyar VJ, Jangid K, Patole MS, Shouche YS (2003) Detection and phylogenetic affiliation of Wolbachia sp. from Indian mosquitoes Culex quinquefasciatus and Aedes albopictus. Current Science 84 (8):1136–1139

