## Supplementary Tables for "Low prevalence of natural *Wolbachia* in major malarial vectors *Anopheles culicifacies* s.l., and *Anopheles stephensi:* a first report"

Table S1: Primers and cycling conditions used for PCR analysis

| **Primer name** | **Sequences** | **Cycling conditions** | **Purpose** |
| --- | --- | --- | --- |
| LCO1490  HCO2198 | 5’-GGTCAACAAATCATAAAGATATTGG  5’-TAAACTTCAGGGTGACCAAAAAATCA | 95° - 2 min,  95° - 30 sec,  46° - 45 sec,  72° - 45 sec,  35 cycles. | Mosquito *COI* amplification |
| WspecF  WspecR | 5’-CATACCTATTCGAAGGGATAG  5’-AGCTTCGAGTGAAACCAATTC | 95° - 2 min,  95° - 30 sec, 55° - 60 sec,  72° - 60 sec,  30 cycles. | *Wolbachia* detection by *16S* rDNA |
| 16SNF  16SNR | 5′-GAAGGGATAGGGTCGGTTCG  5′-CAATTCCCATGGCGTGACG | 95° - 2 min,  95° - 15 sec, 66° - 25 sec,  72° - 30 sec,  35 cycles. | Nested PCR for low density *Wolbachia* detection after the initial amplification by WspecF/R primers |
| Wsp 81F  Wsp 691R | 5′-TGGTCCAATAAGTGATGAAGAAAC  5′-AAAAATTAAACGCTACTCCA | 95° - 2 min,  95° - 15 sec, 66° - 25 sec  72° - 30 sec,  35 cycles. | *Wolbachia* detection by *wsp* gene |
| FtsZ F  FtsZ R | 5’-GTATGCCGATTGCAGAGCTTG  5′-GCCATGAGTATTCACTTGGCT | 95° - 2 min,  95° - 30 sec, 51° - 60 sec,  72° - 60 sec,  35 cycles. | *Wolbachia* detection by *FtsZ* cell cycle gene |
| gatB F1  gatB R1 | 5’-GAKTTAAAYCGYGCAGGBGTT  5’-TGGYAAYTCRGGYAAAGATGA | 94°C - 2 min,  94°C - 30 sec,  54°C - 45 sec,  72°C - 1.5 min,  70°C - 10 min,  36 cycles. | Genotyping *Wolbachia* by glutamyl-tRNA(Gln) amidotransferase, subunit B (*gatB*) |
| coxA F1  coxA R1 | 5’-TTGGRGCRATYAACTTTATAG  5’-CTAAAGACTTTKACRCCAGT | 94°C - 2 min,  94°C - 30 sec,  55°C - 45 sec,  72°C - 1.5 min,  70°C - 10 min,  36 cycles. | Genotyping *Wolbachia* by cytochrome *c* oxidase, subunit I (*CoxA*) |
| hcpA F1  hcpA R1 | 5’-GAAATARCAGTTGCTGCAAA  5’-GAAAGTYRAGCAAGYTCTG | 94°C - 2 min,  94°C - 30 sec,  53°C - 45 sec,  72°C - 1.5 min,  70°C - 10 min,  36 cycles. | Genotyping *Wolbachia* by conserved hypothetical protein (*hcpA)* |
| ftsZ F1  ftsZ R1 | 5’-ATYATGGARCATATAAARGATAG 5’-TCRAGYAATGGATTRGATAT | 94°C - 2 min,  94°C - 30 sec,  54°C - 45 s,  72°C - 1.5 min,  70°C - 10 min,  36 cycles. | Genotyping *Wolbachia* by cell division protein (*ftsZ*) |
| fbpA F1  fbpA R1 | 5’-GCTGCTCCRCTTGGYWTGAT  5’-CCRCCAGARAAAAYYACTATTC | 94°C - 2 min,  94°C - 30 sec,  59°C - 45 sec,  72°C - 1.5 min,  70°C - 10 min,  36 cycles. | Genotyping *Wolbachia* by fructose-bisphosphate aldolase (*fbpA*) |
| Asafor  Asarev | 5’-GCGCGTAGGCGGTTTACAC  5’- AGCGTCAGTAATGAGCCAGGTT | 95º - 2 min,  95º - 30sec,  54º - 60sec,  72º - 60sec,  35 cycles. | *Asaia* detection by targeting *16S rRNA* |

Table S2: Sequences used for *Wolbachia* phylogeny

| **S. No** | **Accession No** | **Source organism** | **Isolate /Strain Id** | **Used sequence** | **Year of isolation** | **Country** |
| --- | --- | --- | --- | --- | --- | --- |
| 1. | ABZA01000001 | *Culex quinquefasciatus* | JHB | Whole genome | 2008 | USA |
| 2. | AF051145 | *Brugia malayi* |  | 16S rRNA partial sequence | 1998 | UK |
| 3. | AF069068 | *Litomosoides sigmodontis* |  | 16S rRNA partial sequence | 1998 | UK |
| 4. | AF397409 | *Culex quinquefasciatus* | NCCS | 16S ribosomal RNA | 2001 | India |
| 5. | AF397410 | *Culex quinquefasciatus* | NIV | 16S ribosomal RNA | 2001 | India |
| 6. | AJ010275 | *Brugia malayi* |  | 16S rRNA partial sequence | 1998 | Italy |
| 7. | AJ010276 | *Onchocerca ochengi* |  | 16S rRNA partial sequence | 1998 | Italy |
| 8. | CAGB01000162 | *Aedes albopictus* | wAlbB | Whole genome | 2011 | France |
| 9. | CP003319 | *Rickettsia massiliae* | AZT80 | Complete genome | 2012 | USA |
| 10. | EU096232 | *Drosophila* | EW-p | 16S rRNA partial sequence | 2007 | South Korea |
| 11. | FR827936 | *Onchocerca skrjabini* | MIB:ZPL:00924 | 16S rRNA partial sequence | 2011 | Italy |
| 12. | FR827944 | *Litomosoides sigmodontis* | MIB:ZPL:01164 | 16S rRNA partial sequence | 2011 | Italy |
| 13. | KJ728740 | *Anopheles gambiae* | VK5_2.9_O | 16S rRNA partial sequence | 2010 | Burkina Faso^ |
| 14. | KJ728741 | *Anopheles gambiae* | VK7_2.3a_O | 16S rRNA partial sequence | 2010 | Burkina Faso^ |
| 15. | KJ728745 | *Anopheles gambiae* | SMS_5.2_O | 16S rRNA partial sequence | 2010 | Burkina Faso^ |
| 16. | KJ728748 | *Anopheles gambiae* | VK7_13.5_O | 16S rRNA partial sequence | 2010 | Burkina Faso^ |
| 17. | KJ728749 | *Anopheles gambiae* | VK7_2.3b_O | 16S rRNA partial sequence | 2010 | Burkina Faso^ |
| 18. | KJ728752 | *Anopheles gambiae* | VK5_7.6_O | 16S rRNA partial sequence | 2010 | Burkina Faso^ |
| 19. | KP089991 | *Anopheles coluzzii* | VK5_8.1.1 | 16S rRNA partial sequence | 2012 | Burkina Faso^‡^ |
| 20. | KU255248 | *Onchocerca eberhardi* | S63-5 244zpl | 16S rRNA partial sequence | 2007 | Japan |
| 21. | KX611381 | *Culex quinquefasciatus* | GC_6b | 16S ribosomal RNA | 2016 | Pakistan |
| 22. | MH596695 | *Anopheles arabiensis* | isolate 13 | 16S rRNA partial sequence | 2018 | Tanzania^#^ |
| 23. | MH596696 | *Anopheles arabiensis* | isolate 15 | 16S rRNA partial sequence | 2018 | Tanzania^#^ |
| 24. | MH596703 | *Anopheles arabiensis* | isolate 140 | 16S rRNA partial sequence | 2018 | Tanzania^#^ |
| 25. | MH605279 | *Anopheles coluzzii* | GHA-DOG1 | 16S rRNA partial sequence | 2017 | Ghana^ø^ |
| 26. | MK026554 | *Aedeomyia madagascarica* | TSA-AMAD-1 | 16S rRNA partial sequence | 2016 | Madagascar |
| 27. | MK026555 | *Culex antennatus* | TSA-CANT-1 | 16S rRNA partial sequence | 2016 | Madagascar |
| 28. | MK026556 | *Culex decens* | TSA-CDEC-1 | 16S rRNA partial sequence | 2016 | Madagascar |
| 29. | MK026557 | *Culex decens* | TSA-CDEC-2 | 16S rRNA partial sequence | 2016 | Madagascar |
| 30. | MK026558 | *Culex duttoni* | TSA-CDUT-1 | 16S rRNA partial sequence | 2016 | Madagascar |
| 31. | MK026559 | *Mansonia uniformis* | TSA-MUNI-1 | 16S rRNA partial sequence | 2016 | Madagascar |
| 32. | MK026560 | *Uranotaenia sp.* | ANI-USP1-1 | 16S rRNA partial sequence | 2016 | Madagascar |
| 33. | MK026561 | *Uranotaenia sp.* | TSA-USP1-1 | 16S rRNA partial sequence | 2016 | Madagascar |
| 34. | MK026562 | *Uranotaenia sp.* | TSA-USP2-1 | 16S rRNA partial sequence | 2016 | Madagascar |
| 35. | MK026563 | *Uranotaenia sp.* | TSA-USP2-2 | 16S rRNA partial sequence | 2016 | Madagascar |
| 36. | MK755768 | *Anopheles carnevalei* | LOP223 | 16S rRNA partial sequence | 2012 | Gabon^$^ |
| 37. | MK755769 | *Anopheles carnevalei* | M39 | 16S rRNA partial sequence | 2012 | Gabon^$^ |
| 38. | MK755770 | *Anopheles coluzzii* | LOP21 | 16S rRNA partial sequence | 2016 | Gabon^$^ |
| 39. | MK755771 | *Anopheles coluzzii* | 553 | 16S rRNA partial sequence | 2016 | Gabon^$^ |
| 40. | MK755772 | *Anopheles coustani* | BNG78 | 16S rRNA partial sequence | 2017 | Gabon^$^ |
| 41. | MK755773 | *Anopheles coustani* | BNG4467 | 16S rRNA partial sequence | 2015 | Gabon^$^ |
| 42. | MK755774 | *Anopheles funestus* | BNG5 | 16S rRNA partial sequence | 2015 | Gabon^$^ |
| 43. | MK755775 | *Anopheles funestus* | BNG6 | 16S rRNA partial sequence | 2015 | Gabon^$^ |
| 44. | MK755784 | *Anopheles gambiae* | BKB5 | 16S rRNA partial sequence | 2014 | Gabon^$^ |
| 45. | MK755785 | *Anopheles gambiae* | BKB6 | 16S rRNA partial sequence | 2014 | Gabon^$^ |
| 46. | MK755786 | *Anopheles gambiae* | BKB7 | 16S rRNA partial sequence | 2014 | Gabon^$^ |
| 47. | MK755787 | *Anopheles hancocki* | BNG2213 | 16S rRNA partial sequence | 2015 | Gabon^$^ |
| 48. | MK755788 | *Anopheles implexus* | M0228 | 16S rRNA partial sequence | 2013 | Gabon^$^ |
| 49. | MK755789 | *Anopheles jebudensis* | B612 | 16S rRNA partial sequence | 2013 | Gabon^$^ |
| 50. | MK755790 | *Anopheles nigeriensis* | BNG3729 | 16S rRNA partial sequence | 2015 | Gabon^$^ |
| 51. | MK755791 | *Anopheles marshallii* | B1197 | 16S rRNA partial sequence | 2013 | Gabon^$^ |
| 52. | MK755792 | *Anopheles marshallii* | BKB17 | 16S rRNA partial sequence | 2014 | Gabon^$^ |
| 53. | MK755793 | *Anopheles moucheti* | B1280 | 16S rRNA partial sequence | 2013 | Gabon^$^ |
| 54. | MK755794 | *Anopheles moucheti* | B1281 | 16S rRNA partial sequence | 2013 | Gabon^$^ |
| 55. | MK755795 | *Anopheles moucheti* | B1282 | 16S rRNA partial sequence | 2013 | Gabon^$^ |
| 56. | MK755823 | *Anopheles nili* | FCV1 | 16S rRNA partial sequence | 2015 | Gabon^$^ |
| 57. | MK755824 | *Anopheles nili* | FCV2 | 16S rRNA partial sequence | 2015 | Gabon^$^ |
| 58. | MK755825 | *Anopheles nili* | FCV5 | 16S rRNA partial sequence | 2015 | Gabon^$^ |
| 59. | MK755834 | *Anopheles paludis* | BTK121 | 16S rRNA partial sequence | 2016 | Gabon^$^ |
| 60. | MK755835 | *Anopheles vinckei* | M002 | 16S rRNA partial sequence | 2012 | Gabon^$^ |
| 61. | MK755836 | *Anopheles vinckei* | M018 | 16S rRNA partial sequence | 2012 | Gabon^$^ |
| 62. | MK755837 | *Anopheles vinckei* | M018 | 16S rRNA partial sequence | 2012 | Gabon^$^ |
| 63. | MN200331 | *Dirofilaria immitis* | Sochi1 | 16S rRNA partial sequence | 2018 | Russia |
| 64. | MN268743 | *Anopheles stephensi* | AE4 | 16S rRNA partial sequence | 2017 | India* |
| 65. | MN268744 | *Anopheles stephensi* | AE5 | 16S rRNA partial sequence | 2017 | India* |
| 66. | MN268746 | *Anopheles stephensi* | AE6 | 16S rRNA partial sequence | 2017 | India* |
| 67. | MN268747 | *Anopheles culicifacies* | TS1 | 16S rRNA partial sequence | 2017 | India* |
| 68. | MN268748 | *Anopheles culicifacies* | TS2 | 16S rRNA partial sequence | 2017 | India* |
| 69. | MN268749 | *Anopheles culicifacies* | TS3 | 16S rRNA partial sequence | 2017 | India* |
| 70. | MN268750 | *Anopheles stephensi* | TS4 | 16S rRNA partial sequence | 2017 | India* |
| 71. | MN880500 | *Anopheles coluzzii* | Anopheles3 | 16S rRNA partial sequence | 2019 | Cameroon |
| 72. | MN880502 | *Anopheles coluzzii* | Anopheles5 | 16S rRNA partial sequence | 2019 | Cameroon |
| 73. | MN887537 | *Anopheles introlatus* | z1 | 16S ribosomal RNA | 2019 | Malaysia^%^ |
| 74. | MN887538 | *Anopheles introlatus* | z4 | 16S ribosomal RNA | 2019 | Malaysia^%^ |
| 75. | MN887539 | *Anopheles introlatus* | Z5 | 16S ribosomal RNA | 2019 | Malaysia^%^ |
| 76. | MN887554 | *Anopheles balabacensis* | m1b3 | 16S ribosomal RNA | 2019 | Malaysia^%^ |
| 77. | MN887555 | *Anopheles balabacensis* | m1b4 | 16S ribosomal RNA | 2019 | Malaysia^%^ |
| 78. | MN887556 | *Anopheles balabacensis* | m1b5 | 16S ribosomal RNA | 2019 | Malaysia^%^ |
| 79. | MN887558 | *Anopheles latens* | m297b | 16S ribosomal RNA | 2019 | Malaysia^%^ |
| 80. | MN887559 | *Anopheles macarthuri* | m296 | 16S ribosomal RNA | 2019 | Malaysia^%^ |
| 81. | MN887560 | *Anopheles sinensis* | SDF1 | 16S ribosomal RNA | 2019 | Malaysia^%^ |
| 82. | MN887561 | *Anopheles sinensis* | SDF1a | 16S ribosomal RNA | 2019 | Malaysia^%^ |
| 83. | MN887562 | *Anopheles sinensis* | SDF3 | 16S ribosomal RNA | 2019 | Malaysia^%^ |
| 84. | MN887564 | *Anopheles hyrcanus* | BLF2 | 16S ribosomal RNA | 2019 | Malaysia^%^ |
| 85. | MN887565 | *Anopheles hyrcanus* | BLF3 | 16S ribosomal RNA | 2019 | Malaysia^%^ |
| 86. | MN887566 | *Anopheles hyrcanus* | BLF4 | 16S ribosomal RNA | 2019 | Malaysia^%^ |
| 87. | MN887573 | *Anopheles barbirostris* | Pu24 | 16S ribosomal RNA | 2019 | Malaysia^%^ |
| 88. | MN887574 | *Anopheles barbirostris* | Tw1 | 16S ribosomal RNA | 2019 | Malaysia^%^ |
| 89. | MN887575 | *Anopheles barbirostris* | Tw2 | 16S ribosomal RNA | 2019 | Malaysia^%^ |
| 90. | MN887581 | *Culex quinquefasciatus* | Pers1 | 16S ribosomal RNA | 2019 | Malaysia |
| 91. | MN887582 | *Culex quinquefasciatus* | Pers7 | 16S ribosomal RNA | 2019 | Malaysia |
| 92. | MN887583 | *Culex quinquefasciatus* | Kla4 | 16S ribosomal RNA | 2019 | Malaysia |
| 93. | MN887584 | *Culex quinquefasciatus* | Kla5 | 16S ribosomal RNA | 2019 | Malaysia |
| 94. | MN887585 | *Culex quinquefasciatus* | Kla29 | 16S ribosomal RNA | 2019 | Malaysia |
| 95. | MT449018 | *Anopheles minimus* | P21 | 16S ribosomal RNA | 2015 | Thailand^∆^ |
| 96. | MT449019 | *Anopheles minimus* | P32 | 16S ribosomal RNA | 2015 | Thailand^∆^ |
| 97. | NC 010981 | *Culex quinquefasciatus* | wPip | Complete genome | 2008 | Srilanka |
| 98. | NC_002978 | *Drosophila melanogaster* | wMel | Complete genome | 2003 | USA |
| 99. | NC_012416 | *Drosophila simulans* | wRi | Complete genome | 2009 | Sweden |
| 100. | NC_021084 | *Drosophila simulans* | wNo | Complete genome | 2012 | Sweden |
| 101. | NR_074459 | *Rickettsia japonica* | YH | 16S rRNA partial sequence | 2013 | USA |
| 102. | NZ_CP021120 | *Chrysomya megacephala* | wMeg | Complete genome | 2015 | Brazil |
| 103. | NZ_CP034334 | *Drosophila mauritiana* | wMau | Complete genome | 2018 | USA |
| 104. | NZ_CP034335 | *Drosophila mauritiana* | wMau | Complete genome | 2018 | USA |
| 105. | NZ_CP041215 | *Carposina sasakii* | wCauA | Complete genome | 2017 | China |
| 106. | NZ_CP042444 | *Drosophila melanogaster* | wMel_I23 | Complete genome | 2019 | USA |
| 107. | NZ_CP042445 | *Drosophila melanogaster* | wMel_ZH26 | Complete genome | 2019 | USA |
| 108. | NZ_CP042446 | *Drosophila melanogaster* | wMel_N25 | Complete genome | 2019 | USA |
| 109. | NZ_CP042904 | *Drosophila ananassae* | W2.1 | Complete genome | 2019 | USA |
| 110. | NZ_CP046577 | *Litomosoides sigmodontis* | wLsig | Complete genome | 2018 | France |
| 111. | NZ_CP046578 | *Dirofilaria immitis* | wDim | Complete genome | 1991 | USA |

* this study, ^ – Baldini et al., (2014), ^#^ – Baldini et al., (2018), ^$^ – Ayala et al., (2019), ^‡^ – Buck et al., (2016), ^%^ – Wong et al., (2020), ^∆^ – Tongkrajang et al., (2020), ^ø^ – Jeffries et al., (2018).

Table S3: Sequences used for *Anopheles* mosquito phylogeny

| **S.No.** | **Accession No.** | **Source organism** | **Isolate/Strain Id** | **Used sequence** | **Year of isolation** | **Country** |
| --- | --- | --- | --- | --- | --- | --- |
| 1. | LR736007 | *Anopheles culicifacies* | TS1 | COI gene, partial cds; mitochondrial | 2017 | India* |
| 2. | LR736008 | *Anopheles culicifacies* | TS2 | COI gene, partial cds; mitochondrial | 2017 | India* |
| 3. | LR736009 | *Anopheles culicifacies* | TS3 | COI gene, partial cds; mitochondrial | 2017 | India* |
| 4. | LR736010 | *Anopheles stephensi* | TS4 | COI gene, partial cds; mitochondrial | 2017 | India* |
| 5. | LR736012 | *Anopheles stephensi* | AE4 | COI gene, partial cds; mitochondrial | 2017 | India* |
| 6. | LR736013 | *Anopheles stephensi* | AE5 | COI gene, partial cds; mitochondrial | 2017 | India* |
| 7. | LR736014 | *Anopheles stephensi* | AE6 | COI gene, partial cds; mitochondrial | 2017 | India* |
| 8. | NC_028223 | *Anopheles stephensi* | ASTEP20150811V3 | Mitochondrion, complete genome | 2015 | USA |
| 9. | MH538704 | *Anopheles stephensi* | voucher BUZOOT | COI gene, partial cds; mitochondrial | 2018 | India |
| 10. | KX467337 | *Anopheles stephensi* | voucher MOSQ02-16 | COX1 gene, partial cds; mitochondrial |  | India |
| 11. | KF406680 | *Anopheles stephensi* | voucher NIBGE DIP-00281 | COI gene, partial cds; mitochondrial | 2007 | Pakistan |
| 12. | NC_028216 | *Anopheles culicifacies* | voucher ACUL20150811V4 | Mitochondrion, complete genome | 2015 | USA |
| 13. | KR732656 | *Anopheles culicifacies* | isolate B | Mitochondrion, complete genome | 2015 | China |
| 14. | KJ010898 | *Anopheles culicifacies* | voucher UNB-04 | CO1 gene, partial cds; mitochondrial | 2013 | UK |
| 15. | DQ424962 | *Anopheles culicifacies* | - | COI) gene, partial cds; mitochondrial | 2006 | India |
| 16. | KR817729 | *Anopheles culicifacies* | voucher BUZOO-M-Ac | COI gene, partial cds; mitochondrial | 2015 | India |
| 17. | KJ010896 | *Anopheles culicifacies* | voucher F04 | COI gene, partial cds; mitochondrial | 2013 | Srilanka |
| 18. | KJ010892 | *Anopheles culicifacies* | voucher H06 | CO1 gene, partial cds; mitochondrial | 2013 | Srilanka |
| 19. | NC_035159 | *Aedes aegypti* | strain LVP_AGWG | Mitochondrion, complete genome | 2017 | USA |

*this study
Table S4: BLAST matches of isolates sequence with reference sequence according to phylogenetic analysis.

| **Query Accession No.** | **Subject Accession No.** | **% Identity** | **Alignment length** | **Mismatches** | **Gap opens** | **Query start** | **Query end** | **Subject start** | **Subject end** | **e-value** | **Bit score** |
| --- | --- | --- | --- | --- | --- | --- | --- | --- | --- | --- | --- |
| MN268747.1 (TS1) | NC_010981.1 | 99.277 | 415 | 0 | 3 | 3 | 417 | 1136966 | 1137377 | 0 | 747 |
|  | NZ_CP021120.1 | 99.277 | 415 | 0 | 3 | 3 | 417 | 597625 | 598036 | 0 | 747 |
|  | NC_021084.1 | 98.795 | 415 | 2 | 3 | 3 | 417 | 704832 | 705243 | 0 | 736 |
|  | NZ_CP034334.1 | 98.795 | 415 | 2 | 3 | 3 | 417 | 198574 | 198985 | 0 | 736 |
|  | NZ_CP034335.1 | 98.795 | 415 | 2 | 3 | 3 | 417 | 198576 | 198987 | 0 | 736 |
|  | CAGB01000162.1 | 98.554 | 415 | 3 | 3 | 3 | 417 | 21802 | 22213 | 0 | 730 |
| MN268748.1 (TS2) | NC_010981.1 | 99.034 | 414 | 2 | 2 | 3 | 416 | 1136966 | 1137377 | 0 | 741 |
|  | NZ_CP021120.1 | 99.034 | 414 | 2 | 2 | 3 | 416 | 597625 | 598036 | 0 | 741 |
|  | NC_021084.1 | 98.551 | 414 | 4 | 2 | 3 | 416 | 704832 | 705243 | 0 | 730 |
|  | NZ_CP034334.1 | 98.551 | 414 | 4 | 2 | 3 | 416 | 198574 | 198985 | 0 | 730 |
|  | NZ_CP034335.1 | 98.551 | 414 | 4 | 2 | 3 | 416 | 198576 | 198987 | 0 | 730 |
|  | CAGB01000162.1 | 98.309 | 414 | 5 | 2 | 3 | 416 | 21802 | 22213 | 0 | 725 |
| MN268749.1 (TS3) | NC_010981.1 | 99.034 | 414 | 2 | 2 | 3 | 416 | 1136966 | 1137377 | 0 | 741 |
|  | NZ_CP021120.1 | 99.034 | 414 | 2 | 2 | 3 | 416 | 597625 | 598036 | 0 | 741 |
|  | NC_021084.1 | 98.551 | 414 | 4 | 2 | 3 | 416 | 704832 | 705243 | 0 | 730 |
|  | NZ_CP034334.1 | 98.551 | 414 | 4 | 2 | 3 | 416 | 198574 | 198985 | 0 | 730 |
|  | NZ_CP034335.1 | 98.551 | 414 | 4 | 2 | 3 | 416 | 198576 | 198987 | 0 | 730 |
|  | CAGB01000162.1 | 98.309 | 414 | 5 | 2 | 3 | 416 | 21802 | 22213 | 0 | 725 |
| MN268750.1 (TS4) | NC_010981.1 | 98.321 | 417 | 5 | 2 | 1 | 417 | 1136963 | 1137377 | 0 | 730 |
|  | NZ_CP021120.1 | 98.321 | 417 | 5 | 2 | 1 | 417 | 597622 | 598036 | 0 | 730 |
|  | NC_021084.1 | 98.321 | 417 | 5 | 2 | 1 | 417 | 704829 | 705243 | 0 | 730 |
|  | NZ_CP034334.1 | 98.321 | 417 | 5 | 2 | 1 | 417 | 198571 | 198985 | 0 | 730 |
|  | NZ_CP034335.1 | 98.321 | 417 | 5 | 2 | 1 | 417 | 198573 | 198987 | 0 | 730 |
|  | CAGB01000162.1 | 99.041 | 417 | 2 | 2 | 1 | 417 | 21799 | 22213 | 0 | 747 |
| MN268743.1 (AE4) | NC_010981.1 | 98.568 | 419 | 2 | 4 | 1 | 419 | 1136963 | 1137377 | 0 | 737 |
|  | NZ_CP021120.1 | 98.568 | 419 | 2 | 4 | 1 | 419 | 597622 | 598036 | 0 | 737 |
|  | NC_021084.1 | 98.091 | 419 | 4 | 4 | 1 | 419 | 704829 | 705243 | 0 | 726 |
|  | NZ_CP034334.1 | 98.091 | 419 | 4 | 4 | 1 | 419 | 198571 | 198985 | 0 | 726 |
|  | NZ_CP034335.1 | 98.091 | 419 | 4 | 4 | 1 | 419 | 198573 | 198987 | 0 | 726 |
|  | CAGB01000162.1 | 97.852 | 419 | 5 | 4 | 1 | 419 | 21799 | 22213 | 0 | 721 |
| MN268744.1 (AE5) | NC_010981.1 | 99.041 | 417 | 2 | 2 | 1 | 417 | 1136963 | 1137377 | 0 | 747 |
|  | NZ_CP021120.1 | 99.041 | 417 | 2 | 2 | 1 | 417 | 597622 | 598036 | 0 | 747 |
|  | NC_021084.1 | 98.561 | 417 | 4 | 2 | 1 | 417 | 704829 | 705243 | 0 | 736 |
|  | NZ_CP034334.1 | 98.561 | 417 | 4 | 2 | 1 | 417 | 198571 | 198985 | 0 | 736 |
|  | NZ_CP034335.1 | 98.561 | 417 | 4 | 2 | 1 | 417 | 198573 | 198987 | 0 | 736 |
|  | CAGB01000162.1 | 98.321 | 417 | 5 | 2 | 1 | 417 | 21799 | 22213 | 0 | 730 |
| MN268746.1 (AE6) | NC_010981.1 | 99.036 | 415 | 1 | 3 | 3 | 417 | 1136966 | 1137377 | 0 | 741 |
|  | NZ_CP021120.1 | 99.036 | 415 | 1 | 3 | 3 | 417 | 597625 | 598036 | 0 | 741 |
|  | NC_021084.1 | 98.554 | 415 | 3 | 3 | 3 | 417 | 704832 | 705243 | 0 | 730 |
|  | NZ_CP034334.1 | 98.554 | 415 | 3 | 3 | 3 | 417 | 198574 | 198985 | 0 | 730 |
|  | NZ_CP034335.1 | 98.554 | 415 | 3 | 3 | 3 | 417 | 198576 | 198987 | 0 | 730 |
|  | CAGB01000162.1 | 98.313 | 415 | 4 | 3 | 3 | 417 | 21802 | 22213 | 0 | 725 |

Table S5: Genetic diversity of *Wolbachia* within groups

|  | **Distance within group (Mean±SE)** |
| --- | --- |
| **Supergroup A** | 0.0012±0.0010 |
| **Supergroup B** | 0.2033±0.2139 |
| **Supergroup C** | 0.0049±0.0041 |
| **Supergroup D** | 0.0025±0.0029 |
| **Outgroup - Rickettsia** | 0.0014±0.0025 |

The number of base substitutions per site from averaging over all sequence pairs within each group is shown above using MCL model. The analysis involved *16S rRNA* gene sequences from 111 *Wolbachia* endosymbionts in *Dipteran* insects. There were a total of 339 positions in the final dataset.

Table S5a: Evolutionary divergence in *Wolbachia* between groups

|  | **Supergroup A** | **Supergroup B** | **Supergroup C** | **Supergroup D** | **Outgroup (Rickettsia)** |
| --- | --- | --- | --- | --- | --- |
| **Supergroup A** |  | **0.1292306** | 0.0099065 | 0.0063562 | 0.042003 |
| **Supergroup B** | **0.1225491** |  | **0.1318681** | **0.1315869** | **0.1583467** |
| **Supergroup C** | 0.0137474 | **0.132629** |  | 0.0110164 | 0.0466666 |
| **Supergroup D** | 0.0078936 | **0.1275433** | 0.0152388 |  |  |
| **Outgroup (Rickettsia)** | 0.0668431 | **0.1757723** | 0.0750448 | 0.0673567 |  |

The number of base substitutions per site from averaging over all sequence pairs between groups is shown. The analysis involved *16S rRNA* gene sequences from 111 *Wolbachia* endosymbionts in *Dipteran* insects. There were a total of 339 positions in the final dataset. Standard error estimate(s) are shown above the diagonal.

Table S6: Evolutionary divergence in *Wolbachia* within and between clades of Supergroup B

|  | **Within Supergroup B**  **(Mean±SE)** | **Between clade of Supergroup B** | | | | | |
| --- | --- | --- | --- | --- | --- | --- | --- |
|  |  | **Clade I** | **Clade II** | **Clade III** | **Clade IV** | **Clade V** | **Clade VI** |
| **Clade_I** | 0.0018±0.0013 |  | 0.003116 | 2.8298191 | 0.0040814 | 0.0030616 | 0.0033768 |
| **Clade_II** | 0.0014±0.0025 | 0.0028285 |  | 2.7952345 | 0.0053374 | 0.0042801 | 0.0046526 |
| **Clade_III** | 0 | 2.5139192 | 2.4909326 |  | 2.8226629 | 2.8206673 | 2.8183249 |
| **Clade_IV** | 0.0028±0.0030 | 0.0046137 | 0.0059316 | 2.5109802 |  | 0.0038512 | 0.0045365 |
| **Clade_V** | 0.0014±0.0016 | 0.0032229 | 0.0045066 | 2.5118204 | 0.0041505 |  | 0.0028089 |
| **Clade_VI** | 0.0007±0.0010 | 0.0034567 | 0.0047506 | 2.5104116 | 0.0653009 | 0.0027446 |  |

The analysis involved *16S rRNA* gene sequences from 111 *Wolbachia* endosymbionts from *Dipteran* insects. There were a total of 339 positions in the final dataset. Standard error estimate(s) are shown above the diagonal for between Clades of Supergroup B.

Table S7: Genetic diversity within *Anopheles* mosquito isolates

| **Isolates** | **Distance within group (Mean±SE)** |
| --- | --- |
| ***Anopheles culicifacies*** | 0.035±0.007 |
| ***Anopheles stephensi*** | 0.052±0.01 |

The number of base substitutions per site from averaging over all sequence pairs within each group are shown using MCL model. The analysis involved 18 *Anopheles* mosquito COI nucleotide sequences. There were a total of 396 positions in the final dataset. Standard error estimate(s) are shown in the last column.

Table S8: Evolutionary divergence between *Anopheles* mosquitos isolates group

| **Isolates** | ***Anopheles culicifacies*** | ***Anopheles stephensi*** |
| --- | --- | --- |
| ***Anopheles culicifacies*** |  | 0.0334994 |
| ***Anopheles stephensi*** | 0.160431 |  |

The number of base substitutions per site from averaging over all sequence pairs between groups are shown using MCL model. The analysis involved 18 *Anopheles* mosquito COI nucleotide sequences. There were a total of 396 positions in the final dataset. Standard error estimate(s) are shown above the diagonal.

Table S9: *Wolbachia* prevalence in *Anopheles* species

| **Sl. No.** | ***Anopheles* species** | **Supergroup** | **Methodology for identification** | **Infection rate (%)** | **Reference** |
| --- | --- | --- | --- | --- | --- |
| 1. | *An. gambiae* | B | *16S rRNA*, *wsp*, *fbpA* PCR; High throughput sequencing *16s rRNA*; shotgun metagenomic sequencing | 11/102 (10.8%) | Baldini et al., (2014) |
|  |  | A | *16S rRNA* PCR, nested PCR & qPCR | 46/69 (67%) | Gomes et al., (2017) |
|  |  | B | *16S rRNA, wsp, ftsZ* PCR; *16S rRNA, ftsZ* RT-PCR | 16/492 (3.2%) | Jeffries et al., (2018) |
|  |  |  | *16S rDNA* nested PCR | 5/44 (11%) | Ayala et al., (2019) |
|  |  | A, B | *16S rDNA*, *wsp*, *CifA*, *CifB*, MLST (*gatB*, *coxA*, *hcpA*, *ftsZ* and *fbpA*) PCR; *16S rRNA* qPCR | 5/286 (1.7%) | Jeffries et al., (2021) |
| 2. | *An. coluzzii* | Arthropod-specific supergroup | *16S rRNA*, *wsp*, *fbpA* PCR; High throughput sequencing *16s rRNA*; shotgun metagenomic sequencing | 10/78 (12.8%) | Baldini et al., (2014) |
|  |  | Arthropod-specific supergroup | *16S* rDNA, *16S rDNA* nested PCR | 275/602 (46%) | Shaw et al., (2016) |
|  |  | B | *16S rRNA, wsp, ftsZ* PCR; *16S rRNA, ftsZ* RT-PCR | 12/287 (4.2%) | Jeffries et al., (2018) |
|  |  | A, B | *16S rDNA* nested PCR | 2/58 (3-4%) | Ayala et al. (2019) |
|  |  | B | *16S rRNA*, *16S rRNA* nested PCR and sequencing | 102/171 (59.6%) | Straub et al., (2020) |
| 3. | *An. arabiensis* | A, B | *16S rRNA, wsp, ftsZ* PCR; *16S rRNA, ftsZ* RT-PCR | 1/18 (5.55%) | Jeffries et al., (2018) |
|  |  |  | *16S rDNA* nested PCR | 13/212 (6.1%) | Baldini et al., (2018) |
| 4. | *An. moucheti* | B | *16S rRNA, wsp, ftsZ* PCR; *16S rRNA, ftsZ* RT-PCR | 1/1 (100%) | Jeffries et al., (2018) |
|  |  |  | *16S rDNA* nested PCR | 30/42 (71%) | Ayala et al., (2019) |
|  |  |  | *16S rRNA* RT-PCR; FISH | 621/1093 (56.8%); Ovary (FISH) - 9/16 (56.3%) | Walker et al., (2020) |
| 5. | *An. species A*  *(An. demeilloni)* | B | *16S rRNA, wsp, ftsZ* PCR; *16S rRNA, ftsZ* RT-PCR | 31/35 (88.5%) | Jeffries et al., (2018) |
|  |  |  | *16S rDNA*, *wsp*, *CifA*, *CifB*, MLST (*gatB*, *coxA*, *hcpA*, *ftsZ* and *fbpA*) PCR | 1/1 (100%) | Jeffries et al., (2019) |
|  |  |  | *16S rRNA* RT-PCR; FISH | 284/488 (58.2%) | Walker et al., (2020) |
|  |  |  | *16S rDNA*, *wsp*, *CifA*, *CifB*, MLST (*gatB*, *coxA*, *hcpA*, *ftsZ* and *fbpA*) PCR; *16S rRNA* qPCR | 1/1 (100%) | Jeffries et al., (2021) |
| 6. | *An. funestus* | A, B | *16S rDNA* PCR, nested PCR;  Confirmation by qPCR | 3/247 (1.2%) | Niang et al., (2018) |
|  |  | A | *16S rDNA* nested PCR | 2/37 (5%) | Ayala et al. (2019) |
| 7. | *An. melas* | A | *16S rDNA*, *wsp*, *CifA*, *CifB*, MLST (*gatB*, *coxA*, *hcpA*, *ftsZ* and *fbpA*) | 18/168 (10.7%) | Jeffries et al., (2019) |
|  |  | A, B | *16S rDNA*, *wsp*, *CifA*, *CifB*, MLST (*gatB*, *coxA*, *hcpA*, *ftsZ* and *fbpA*) PCR; *16S rRNA* qPCR | 16/140 (11.4%) | Jeffries et al., (2021) |
| 8. | *An. carnevalei* | A, B | *16S rDNA* nested PCR | 2/29 (7%) | Ayala et al., (2019) |
| 9. | *An. coustani* | B, C* | *16S rDNA* nested PCR | 2/35 (6%) | Ayala et al. (2019) |
| 10. | *An. hancocki* | B | *16S rDNA* nested PCR | 1/41 (2%) | Ayala et al., (2019) |
| 11. | *An. implexus* | B | *16S rDNA* nested PCR | 1/26 (4%) | Ayala et al., (2019) |
| 12. | *An. jebudensis* | B | *16S rDNA* nested PCR | 1/2 (50%) | Ayala et al., (2019) |
| 13. | *An. marshallii* | B | *16S rDNA* nested PCR | 2/42 (5%) | Ayala et al., (2019) |
| 14. | *An. nigeriensis* | B | *16S rDNA* nested PCR | 1/27 (4%) | Ayala et al., (2019) |
| 15. | *An. nili* | B | *16S rDNA* nested PCR | 11/19 (58%) | Ayala et al., (2019) |
| 16. | *An. paludis* | B | *16S rDNA* nested PCR | 1/16 (6%) | Ayala et al., (2019) |
| 17. | *An. vinckei* | A, B | *16S rDNA* nested PCR | 3/30 (10%) | Ayala et al., (2019) |
| 18. | *An. minimus* | D, F | *16S rDNA* nested PCR, qPCR | 4/90 (4.5%) | Sawasdichai et al., (2019) |
| 19. | *An. maculatus* | B, F | *16S rDNA* nested PCR, qPCR | 4/90 (4.5%) | Sawasdichai et al., (2019) |
|  |  | B | *wsp*, *16S rRNA* nested PCR | 2/9 (22.2%) | Wong et al., (2020) |
| 20. | *An. pseudowillmori* | B | *16S rDNA* nested PCR, qPCR | 1/11 (9%) | Sawasdichai et al., (2019) |
| 21. | *An. sawadwongporni* | B | *16S rDNA* nested PCR, qPCR | 1/68 (1.5%) | Sawasdichai et al., (2019) |
| 22. | *An. baimaii* | B, D | *16S rDNA* nested PCR, qPCR | 2/99 (2%) | Sawasdichai et al., (2019) |
| 23. | *An. dirus* | B | *16S rDNA* nested PCR, qPCR | 1/12 (8%) | Sawasdichai et al., (2019) |
| 24. | *An. balabacensis* | B | *wsp*, *16S rRNA* nested PCR | 4/19 (21%) | Wong et al., (2020) |
| 25. | *An. latens* | A, B | *wsp*, *16S rRNA* nested PCR | 4/8 (50%) | Wong et al., (2020) |
| 26. | *An. introlatus* | A, B | *wsp*, *16S rRNA* nested PCR | 17/54 (31.5%) | Wong et al., (2020) |
| 27. | *An. macarthuri* | B | *wsp*, *16S rRNA* nested PCR | 1/4 (25%) | Wong et al., (2020) |
| 28. | *An. barbirostris* | B | *wsp*, *16S rRNA* nested PCR | 5/10 (50%) | Wong et al., (2020) |
| 29. | *An. hyrcanus* | B | *wsp*, *16S rRNA* nested PCR | 9/18 (50%) | Wong et al., (2020) |
| 30. | *An. sinensis* | B | *wsp*, *16S rRNA* nested PCR | 4/7 (57.1%) | Wong et al., (2020) |

Supergroup C – found only in filarial worms. *C – Filarial nematode - *Dirofilaria immitis* was observed in the specimen (Ayala et al 2019).

**Reference:**

Ayala D, Akone-Ella O, Rahola N, Kengne P, Ngangue MF, et al. 2019. Natural Wolbachia infections are common in the major malaria vectors in Central Africa. *Evolutionary Applications* 12: 1583-94

Baldini F, Rouge J, Kreppel K, Mkandawile G, Mapua SA, et al. 2018. First report of natural Wolbachia infection in the malaria mosquito Anopheles arabiensis in Tanzania. *Parasites & vectors* 11: 635

Baldini F, Segata N, Pompon J, Marcenac P, Shaw WR, et al. 2014. Evidence of natural Wolbachia infections in field populations of Anopheles gambiae. *Nature communications* 5: 3985

Buck M, Nilsson LK, Brunius C, Dabiré RK, Hopkins R, Terenius O. 2016. Bacterial associations reveal spatial population dynamics in Anopheles gambiae mosquitoes. *Sci Rep* 6: 22806

Gomes FM, Hixson BL, Tyner MDW, Ramirez JL, Canepa GE, et al. 2017. Effect of naturally occurring Wolbachia in Anopheles gambiae s.l. mosquitoes from Mali on Plasmodium falciparum malaria transmission. *Proceedings of the National Academy of Sciences of the United States of America* 114: 12566-71

Jeffries CL, Cansado-Utrilla C, Beavogui AH, Stica C, Lama EK, et al. 2021. Evidence for natural hybridization and novel Wolbachia strain superinfections in the Anopheles gambiae complex from Guinea. *R. Soc. open sci.* 8

Jeffries CL, Cansado-Utrilla C, Stica C, Walker T. 2019. High density Novel *Wolbachia* strains in *Anopheles* species from Guinea. *bioRxiv*: 772855

Jeffries CL, Lawrence GG, Golovko G, Kristan M, Orsborne J, et al. 2018. Novel Wolbachia strains in Anopheles malaria vectors from Sub-Saharan Africa. *Wellcome open research* 3: 113

Niang EHA, Bassene H, Makoundou P, Fenollar F, Weill M, Mediannikov O. 2018. First report of natural Wolbachia infection in wild Anopheles funestus population in Senegal. *Malaria journal* 17: 408

Sawasdichai S, Chaumeau V, Dah T, Kulabkeeree T, Kajeechiwa L, et al. 2019. Detection of diverse Wolbachia 16S rRNA sequences at low titers from malaria vectors in Kayin state, Myanmar. *Wellcome open research* 4: 11

Shaw WR, Marcenac P, Childs LM, Buckee CO, Baldini F, et al. 2016. Wolbachia infections in natural Anopheles populations affect egg laying and negatively correlate with Plasmodium development. *Nature communications* 7: 11772

Straub TJ, Shaw WR, Marcenac P, Sawadogo SP, Dabire RK, et al. 2020. The Anopheles coluzzii microbiome and its interaction with the intracellular parasite Wolbachia. *Sci Rep* 10: 13847

Tongkrajang N, Ruenchit P, Tananchai C, Chareonviriyaphap T, Kulkeaw K. 2020. Molecular identification of native Wolbachia pipientis in Anopheles minimus in a low-malaria transmission area of Umphang Valley along the Thailand-Myanmar border. *Parasites & vectors* 13: 579

Walker T, Quek S, Jeffries CL, Bandibabone J, Dhokiya V, et al. 2020. Genomic and microscopic evidence of stable high density and maternally inherited *Wolbachia* infections in *Anopheles* mosquitoes. *bioRxiv*: 2020.10.29.357400

Wong ML, Liew JWK, Wong WK, Pramasivan S, Mohamed Hassan N, et al. 2020. Natural Wolbachia infection in field-collected Anopheles and other mosquito species from Malaysia. *Parasites & vectors* 13: 414
